# Refinement of the Pathogenic T Cell Receptor Motif Suggests Type 17 CD8+ T Cells Initiate HLA-B*27-associated Autoimmunity

**DOI:** 10.64898/2026.09.18.752092

**Authors:** Isabel Risch, Qingping Wu, Rahul Devkota, Sophia Y. Li, Joy Um, Tertuliano Alves Pereira Neto, Sarah Walden, Paulo Henrique Arantes de Faria, Andrei Belean, Lucy Durham, Michael Tang, Zoya Qaiyum, Robert D. Inman, Leonie S. Taams, Xinbo Yang, Philip A. Mudd, Nicholas C. Borcherding, Malachi Griffith, Obi L. Griffith, Michael A. Paley

## Abstract

Human leukocyte antigen B*27 (HLA-B*27) is a major risk factor for autoimmune diseases such as axial spondyloarthritis (axSpA), acute anterior uveitis (AAU), and psoriatic arthritis (PsA). HLA-B*27 presents bacterial and self-antigens to disease-associated CD8+ T cells, however, how this leads to a pathogenic IL-17-mediated disease remains unclear. We performed mutational experiments in an established disease-associated TCR and demonstrated that the pathogenic motif encompasses a wider range of TCR sequences than previously appreciated. We next leveraged computational pairing of alpha/beta TCR sequencing (TCR-seq) to demonstrate that the broadened pathogenic motif distinguishes axSpA and AAU patients from healthy controls. Paired single-cell RNA sequencing and single-cell TCR sequencing datasets demonstrated that the pathogenic cells have a distinct transcriptional signature. We find a coupling of pathogenic TCRs to this transcriptional signature in public datasets from axSpA and PsA joint fluid and AAU ocular fluid. Finally, we detect the pathogenic CD8+ T cells in the gut and demonstrate that they express a Type 17 program. Our findings suggest that HLA-B*27+ axSpA, AAU, and PsA are initiated by HLA-B*27-restricted pathogenic CD8+ T cells that undergo Type 17 differentiation in response to intestinal antigens.

**One Sentence Summary:** Multiple HLA-B*27-associated autoimmune diseases share the same pathogenic CD8+ T cells with Type 17 gene expression and cytokine production.

## INTRODUCTION

Human leukocyte antigen B*27 (HLA-B*27) is a major risk factor for autoimmune diseases such as axial spondyloarthritis (axSpA), acute anterior uveitis (AAU), reactive arthritis, enteropathic arthritis, and psoriatic arthritis (PsA), but the exact mechanism of pathogenesis is not fully understood(*1–6*). HLA-B*27 appears to promote autoimmunity by activating antigen-specific CD8+ T cells(*2, 7*). A recurring “public” T cell receptor (TCR) motif has been defined in CD8+ T cells across HLA-B*27+ axSpA, AAU, and reactive arthritis(*2, 7–9*). This TCRꞵ motif consists of the *TRBV9* gene segment and a CDR3ꞵ sequence of (C-A-S-S-[V/L/P]-[G/A]-[L/T]-[Y/F]-S-T-D-T-Q)(*2*). High-throughput TCRꞵ sequencing has demonstrated that this motif is enriched in HLA-B*27+ patients over HLA-B*27+ healthy controls(*7, 10*). These TCRꞵ features are paired with the TRAV21 gene segment in the TCRɑ. CD8+ T cells with these TCR motifs are expanded in inflamed tissues (joint in axSpA, eye in AAU) over peripheral blood. Moreover, in a case study and a subsequent phase II trial, depletion via an anti-TRBV9 monoclonal antibody (seniprutug) was able to ameliorate disease, providing proof-of-concept that these cells are pathogenic(*11*).

The identification of a public TCR motif that is shared across patients and tissues suggested that HLA-B*27 presents a specific peptide (or set of peptides) to CD8+T cells. A yeast-based library screen identified cognate antigens and found that the public TCR motif cross-reacts to the bacterial antigen YeiH as well as a short list of human autoantigens(*7*). The YeiH antigen is derived from Gram-negative intestinal bacteria, such as *E. coli, Salmonella,* and *Shigella*(*7*). Notably, *Salmonella* and *Shigella* are species known to trigger reactive arthritis, suggesting that the initial activation of these CD8+ T cells occurs in intestinal tissues(*6*). Consistent with this hypothesis, enteropathic arthritis is characterized by comorbid inflammatory bowel disease (IBD) and joint involvement, and axSpA patients have been shown to display subclinical levels of gut inflammation(*12, 13*). The accumulated evidence suggests a model where pathogenic CD8+ T cells are first activated by bacterial antigens like YeiH in the gut and subsequently react to structurally similar self-antigens in the joint and the eye.

An increasing body of evidence also suggests that Type 17 responses are central to the pathogenesis of spondyloarthropathy. PsA and axSpA patients display robust responses to IL-17 inhibitors such as secukinumab, ixekizumab, and bimekizumab(*14, 15*). CD4+ T cells, namely T helper 17 (Th17) cells, are the most well-recognized producers of Type 17 cytokines and are thought to be the main cell type promoting the Type 17 response in the joint(*16, 17*). Notably, other cell types also may produce Type 17 cytokines in axSpA, including mucosal-associated invariant T (MAIT) cells(*18*). How the antigen-specific activation of CD8+ T cells, which are traditionally thought to be cytotoxic, leads to a Type 17 immune response led by CD4+ T cells remains unresolved.

In PsA, CD8+ T cells have been recognized as potential contributors to Type 17 cytokine production(*19*), however, whether this is in response to autoantigens remains unclear. By contrast, Type 17 CD8+ T cells (Tc17s) have remained under-studied in axSpA. Pathogenic CD8+ T cells express intermediate levels of CD161(*9*), a marker for IL-17-producing T cell subsets(*20*), suggesting that CD8+ T cells may be a key initiator of the pathogenic IL-17 response in axSpA.

Here, we perform mutagenesis experiments to expand our understanding of the pathogenic TCR motif. We then use that expanded motif to investigate the pathogenic cells in single-cell RNA sequencing (scRNA-seq) and single-cell TCR sequencing (scTCR-seq) datasets. We show that the pathogenic CD8+ T cells are characterized by a distinct gene signature, are present in the gut of HLA-B*27+ individuals, express canonical Type 17 genes, and produce IL-17 in response to *in vitro* stimulation. Taken together, our findings suggest that spondyloarthritis begins in the gut with an anti-bacterial, Type 17 CD8+ T cell response which is brought to the joint and the eye due to cross-reactivity to autoantigens.

## RESULTS

### Specific TCR features allow CD8+ T cell cross-reactivity in axSpA and AAU

Reported pathogenic TCRs in axSpA and AAU often use *TRBV9* for the TRBV gene segment, however, other TRBVs (namely *TRBV5-5*) have been reported(*9*). Pathogenic TCRs recognize the bacterial antigen YeiH and several human autoantigens (GPER1, PRPF3, and RNASEH2B) via the CDR3β loop and the CDR1*a* loop (encoded by *TRAV21*), while the CDR2α and CDR2ꞵ contact HLA-B*27(*7, 9*) (**Fig. 1A**). However, the total scope of TCRs that contain this cross-reactive property has not been defined. We sought to broaden our understanding of the “permitted” pathogenic TCRβ sequences via a series of saturation mutagenesis experiments in the TCRβ chain. We used the UV180.2 TCR as a template, as it is an intraocular, clonally expanded TCR from AAU which has been previously shown to cross-react to the YeiH.232-240 (YeiH) peptide as well as autoantigens(*7, 9*). We made systematic alterations to the TRBV gene and the CDR3β motif at positions 5-10 (**Fig. 1B**). We then transduced TCR-deficient Jurkat cells containing an NFAT-GFP reporter with the altered TCRs and assessed their ability to react to YeiH and the human autoantigens GPER1, PRPF3, and RNASEH2B, presented on HLA-B*27+ K562 cells(*7, 9, 21*) (**Supplementary Materials and Methods**).

**Fig. 1.**
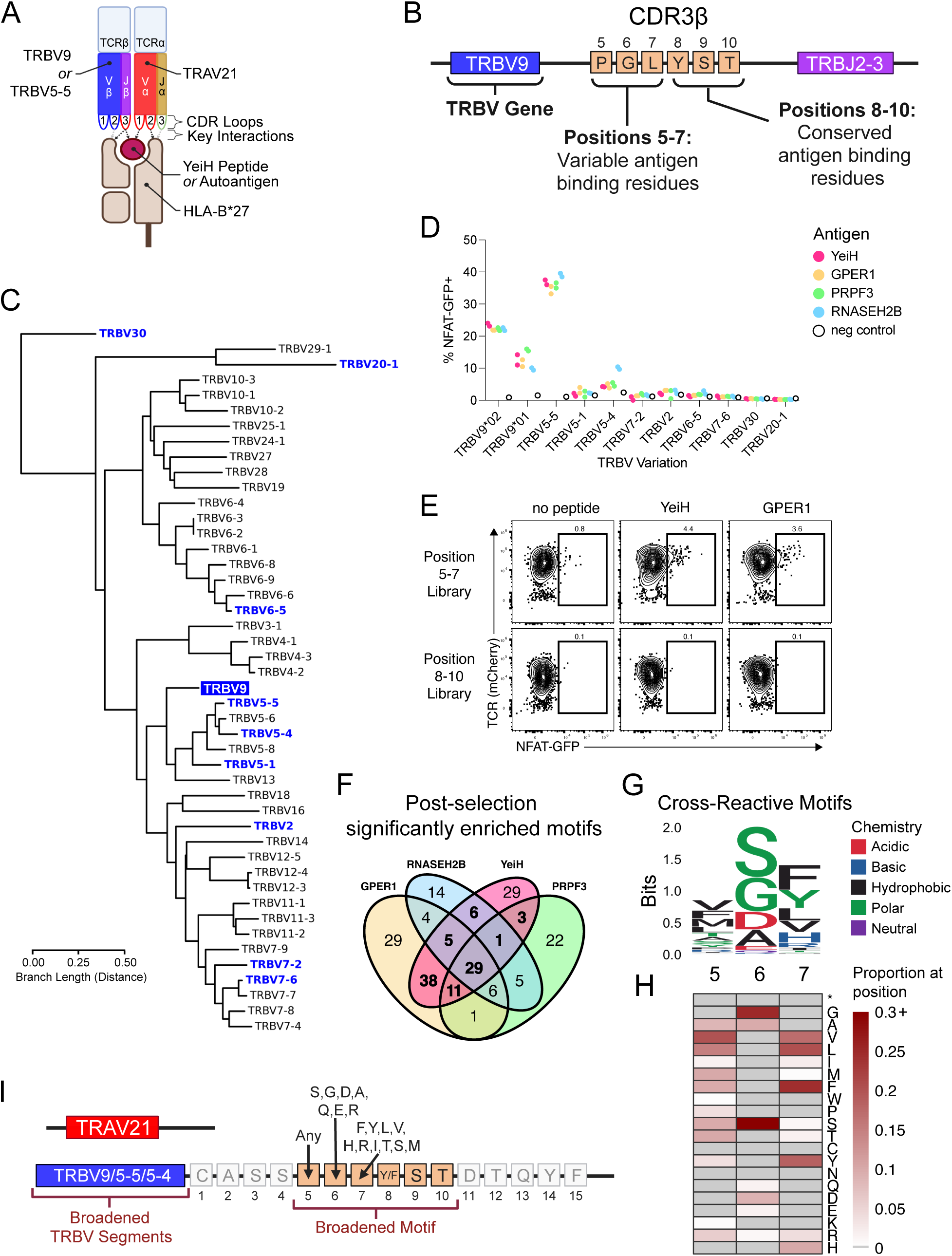
Mutational analysis reveals a broadened TCRꞵ motif for pathogenic CD8+ T cells in axSpA and AAU. **(A)** Illustration of key interactions between the HLA-B*27-antigen complex (bottom) and a pathogenic TCR (top). Black arrows indicate strong interactions; gray arrows indicate weak interactions. **(B)** Illustration of the native TCRꞵ in UV180.2 with regions altered in the mutational experiments. **(C)** Distance tree of TRBV gene segments. TRBV9 is highlighted in blue. TRBV substitutions are shown in blue text. **(D)** Reactivity to the bacterial antigen YeiH and indicated autoantigens (GPER1, PRPF3, and RNASEH2B) was measured using an NFAT-GFP reporter for the indicated TRBV substitutions of *TRBV9*02*. **(E)** Representative flow cytometry from two CDR3ꞵ mutational libraries after exposure to no peptide, YeiH, or GPER1. Top row, position 5-7 library; bottom row, position 8-10 library. **(F)** Venn diagram of the enriched motifs in the 5-7 library after exposure to YeiH and indicated autoantigens. Cross-reactive motifs are shown in bold. **(G** and **H)** Cross-reactive motifs from (F) are summarized as a SeqLogo diagram (G) or a heatmap for at each position (H). Grey boxes in (H) indicate amino acids that were not detected; asterisk (*) indicates a STOP codon. **(I)** Illustration of the broadened pathogenic TCR motif based on panels (G) and (H).

In our first mutational approach, we substituted a panel of candidate TRBV gene segments to replace *TRBV9*02,* the native TRBV gene of UV180.2. We selected 10 TRBV gene segments with a range of amino acid sequence similarities(*22, 23*) (**Fig. 1C**). The candidate gene segments included *TRBV9*01,* the other allele of *TRBV9*; *TRBV5-5*, most similar TRBV gene to *TRBV9* in the UniProt database; *TRBV20-1,* the most dissimilar TRBV gene to *TRBV9* in the UniProt database; and 7 gene segments in between (*TRBV2, TRBV5-1, TRBV5-4, TRBV6-5, TRBV7-2, TRBV7-6,* and *TRBV30*)(*24*). We found that *TRBV9*02*, *TRBV9*01,* and *TRBV5-5* supported a robust response to all peptides, while substitution with *TRBV5-4* led to a more modest signal (**Fig. 1D**). *TRBV5-1* was non-reactive to all antigens despite being the second most similar sequence to *TRBV9* in our panel. Our findings suggest that *TRBV9* and *TRBV5-5* may be the most supportive TRBV gene segments for cross-reactivity, followed by *TRBV5-4*.

Next, we performed a library screen on positions 5-10 of the CDR3ꞵ motif within UV180.2. For our screen, we divided the CDR3ꞵ into two separate mutational analyses to limit each library to less than 40,000 variants for practical execution. In one library, we mutated the “P-G-L” at positions 5-7, while in the second library, we mutated the “Y-S-T” motif at positions 8-10. For both cases, we transduced the TCR library into Jurkat cells and measured the amount of NFAT-GFP reporter activity in response to peptide stimulation. In the 5-7 library, we found a small but consistent population responding to YeiH and autoantigens (**Fig. 1E**). However, we did not detect any NFAT-GFP reporter activity above background for positions 8-10, demonstrating that alteration of this core motif abolished TCR signaling for >99% of possible sequences and consistent with the key role of the “Y-S-T” sequence in peptide recognition. Therefore, we focused our downstream analysis on the position 5-7 library.

For positions 5-7, we sorted the NFAT-GFP+ responders and sequenced their CDR3ꞵ to compare the pre- and post-selection libraries. Across all antigens, 82% of motifs from the initial library were lost after sorting on GFP+ cells (**Fig. S1, A to D**). We defined 93 motifs enriched after YeiH stimulation and enriched in one or more human autoantigen conditions as “cross-reactive” (**Fig. 1F**). Consistent with published *in vivo* data, position 5 appeared minimally conserved in the cross-reactive motifs with 13/20 possible amino acids represented, while position 6 showed the highest level of conservation(*8*) (**Fig. 1G**). As expected, position 6 showed overrepresentation of glycine (25%) and alanine (10%), although serine was the most highly represented (52%) and 4 other amino acids were detected at low (< 10%) levels in the position(*7, 8*) (**Fig. 1G-1H**). In position 7, 10/20 possible amino acids were detected; phenylalanine made up the highest percentage of the results (24%), followed by leucine (20%), tyrosine (19%) and valine (17%). There appeared to exist no strong patterns of linkage between adjacent amino acids (**Fig. S1E**). Our results recapitulate reported sequences from literature while also providing evidence for an expansion of the canonical pathogenic CDR3ꞵ motif in axSpA and AAU. Although the [Y/F]-S-T motif in positions 8-10 appears essential for reactivity, the pre-YST motif (positions 5-7) appears more flexible than previously assumed, with the strongest conservation at position 6. Based on these data, we expanded our definition of the pathogenic CDR3ꞵ motif as follows: position 5 is unrestricted, positions 6 and 7 must contain an amino acid detected at those positions in our screen, and positions 8-10 must contain the canonical [Y/F]-S-T (**Fig. 1I**).

### Broadened TCR motif is increased in patients versus healthy controls

We next asked whether we could use the broadened TCRꞵ motif to better detect pathogenic TCRs in the blood in order to distinguish HLA-B*27+ healthy individuals from HLA-B*27+ patients with axSpA and/or AAU(*7*). We developed three progressively stringent approaches for calling pathogenic TCRs (Criteria 1, 2, and 3), all based on the broadened motif. In brief, Criteria 1 only required the CDR3ꞵ sequence to match the broadened motif; Criteria 2 required both the CDR3ꞵ sequence and the *TRBV* gene segment to match the motif; and Criteria 3 required the CDR3ꞵ, the *TRBV* gene segment, and the *TRAV* gene segment to match (see **Supplementary Materials and Methods**) (**Fig. 2A**). We then applied these criteria to high-throughput TCRseq data and calculated the proportions of the defined motifs in each sample. Using these proportions, we evaluated each test’s ability to distinguish healthy controls from patients. For our approach, we chose to use throughput-intensive rapid TCR library sequencing (TIRTL-seq), a recently published, open-source platform for high-throughput paired TCR sequencing(*25*). We performed TIRTL-seq on a cohort of 8 HLA-B*27+ healthy controls and 6 HLA-B*27+ patients (**Data File S1**). TIRTL-seq returned an average of 57,234,124 alpha and 94,461,307 beta chain reads per sample (**Fig. S2A**), which were resolved into an average of 725,554 unique TCRꞵ chain sequences and 652,028 unique TCRα chain sequences per sample (**Fig. 2B**). Computational pairing led to an average of 39,845 unique full αꞵTCRs per sample.

**Fig. 2.**
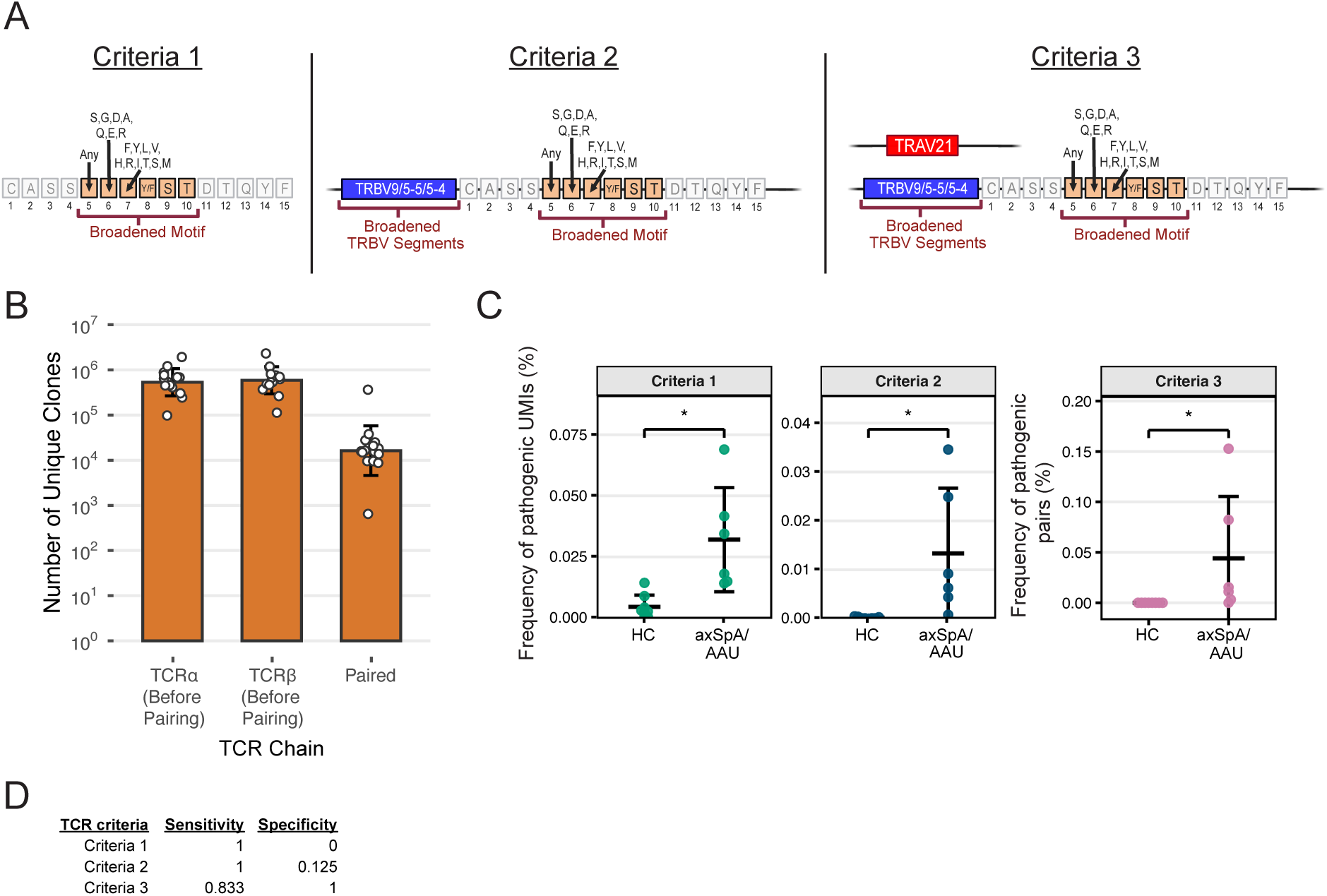
Broadened pathogenic TCR motif distinguishes axSpA patients from healthy controls. **(A)** Illustration of the three pathogenic criteria used to distinguish patients from healthy controls. “Criteria 1” uses the CDR3ꞵ motif, “Criteria 2” uses CDR3ꞵ and TRBV, and “Criteria 3” uses CDR3ꞵ, TRBV, and paired TRAV. **(B)** Barplots of the number of unique TCRα, TCRꞵ, and paired alpha-beta TCR sequences detected using TIRTL-seq. Bars indicate geometric means; error bars are geometric standard deviation. *n =* 14 TIRTL-seq samples. **(C)** Testing Criteria 1-3 to distinguish healthy controls from patients in TIRTLseq. Error bars indicate the mean and standard deviation. **(D)** Sensitivity and specificity tables for (C).

We next assessed the capacity of Criteria 1-3 to distinguish healthy individuals from axSpA and/or AAU (**Fig. 2C**). As paired alpha-beta sequences made up a subset of the total alpha and beta reads, we applied Criteria 1-2 to all TCRꞵ reads and Criteria 3 to the paired alpha-beta reads, to maximize detection of the target motif. All three criteria significantly separated patients from healthy controls (**Fig. 2C**). We next asked how well the criteria distinguish patients from healthy controls based on the presence or absence of the target TCR motif. We found that Criteria 2 was able to identify axSpA/AAU with 100% sensitivity (ability to correctly detect true patients) in this small cohort, however, specificity (ability to correctly call true healthy controls) was limited (12.5%). Criteria 3, meanwhile, achieved 83% sensitivity and 100% specificity (**Fig. 2D**). These results provide proof-of-concept that a search for pathogenic TCR features, particularly one incorporating both alpha- and beta-chain features, can discriminate between healthy controls and patients using peripheral blood alone.

### Pathogenic CD8+ T cells express a distinct transcriptional program compared to other YeiH-specific CD8+ T cells

We next sought to characterize the population of pathogenic CD8+ T cells that contribute to the YeiH-specific response in HLA-B*27+ healthy controls and participants with axSpA and/or AAU. We isolated CD8+ T cells from the peripheral blood of five HLA-B*27+ healthy controls and four HLA-B*27+ AAU patients. To obtain sufficient cells for sequencing, we selected samples that were in the top 25% of YeiH+ CD8+ T cell frequency. Of the AAU individuals, two had AAU alone and two had both AAU and axSpA (**Data File S2)**. We sorted YeiH+ CD8+ T cells and performed paired scRNA-seq and scTCR-seq (**Fig. 3A**).

**Fig. 3.**
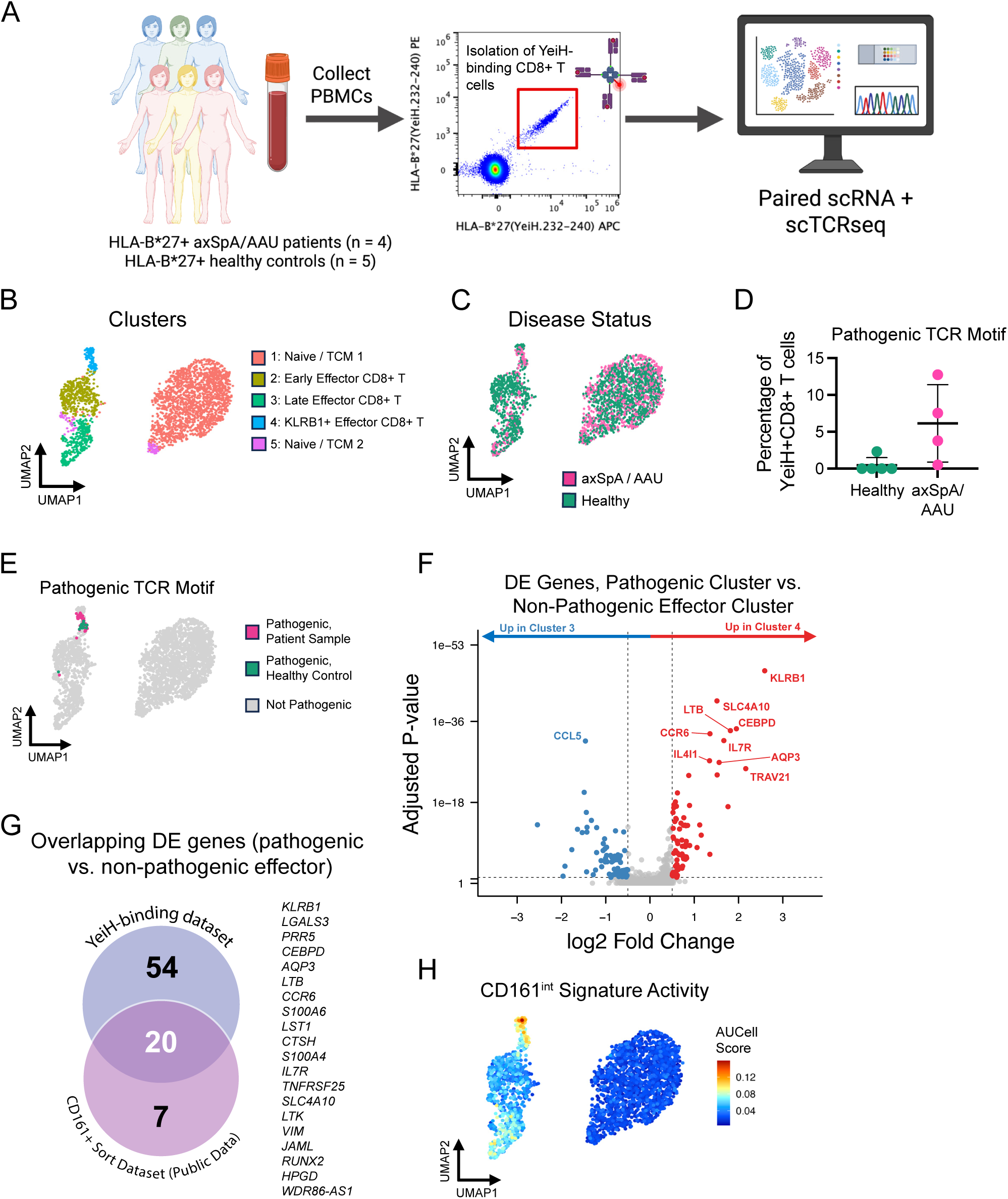
Pathogenic CD8+ T cells express a distinct transcriptional program compared to other YeiH-specific CD8+ T cells. **(A)** Schematic of YeiH+CD8+ T cell isolation from the peripheral blood of HLA-B*27+ healthy controls and patients with axSpA and/or AAU. **(B)** UMAP displaying Seurat clustering of YeiH-sorted CD8+ T cells (2,652 cells total). **(C)** UMAP colored by disease status. **(D)** Percentage of YeiH+CD8+ T cells in each sample which contain pathogenic TCR sequences. **(E)** UMAP plot highlighting the cells with historical pathogenic TCRs. Pathogenic cells are colored based on whether they come from an AxSpA / AAU patient (pink) or a healthy control (green). **(F)** Volcanoplot displaying DEGs in the pathogenic cell-containing cluster (Cluster 4) versus the non-pathogenic effector cluster (Cluster 3). Red color indicates significantly (padj < 0.05 and log2FC > 0.5) higher expression in Cluster 4; blue color indicates significantly higher expression in Cluster 3. The 10 most significant DEGs by adjusted p-value are labeled. **(G)** Venn diagram summarizing the overlap of the DEGs from panel (H) (top) versus the previously published CD161+ sort dataset (bottom). Genes at right are the 20 genes included in the signature. **(H)** CD161int signature activity, calculated using the AUCell algorithm.

Unsupervised clustering of the scRNA-seq data revealed five distinct clusters (**Fig. 3B**). There were no significant differences in cluster distribution between healthy participants and participants with axSpA or AAU (**Fig. 3C, Fig. S3, A to B**). Cluster 3 displayed high activity of cytotoxic molecules and an effector CD8+ T cell signature(*26, 27*), suggesting a “late” or terminally differentiated effector phenotype **(Fig. S3, C to D**). Cluster 4 also expressed the effector signature alongside a number of unique genes such as *KLRB1*, *CEBPD,* and *SLC4A10* (**Fig. S3C).** After clustering the gene expression data, we examined the paired single-cell TCR sequences. Most of Cluster 1 (naive/CM-like) were singlets. By contrast, the effector clusters showed evidence of clonal expansion (**Fig. S3, E to G**). We used Criteria 3 from **Fig. 2A** as our criteria to detect pathogenic TCRs and found that 54 cells total (2% of the dataset) contained pathogenic TCRs. We detected pathogenic CD8+ T cells in all four participants with axSpA and/or AAU (range: 1 - 19 cells per sample) **(Fig. 3D).** 14 pathogenic cells were also detected in one of the five healthy controls, suggesting that autoreactive CD8+ T cells may arise in some healthy individuals, similar to Type 1 diabetes(*28*). Notably, the frequency of pathogenic CD8+ T cells was greater in three of the four axSpA/AAU participants compared to the healthy control **(Fig. 3D),** suggesting expansion of pathogenic CD8+ T cells may facilitate development of AAU and axSpA. We next mapped the pathogenic TCRs to the single-cell RNA expression and found that 49/54 (91%) of the pathogenic cells localized to the *KLRB1*^+^ effector cluster (Cluster 4; **Fig. 3E**). Collectively, these data suggest that pathogenic CD8+ T cells are a small fraction of the total YeiH-specific response and are transcriptionally distinct from most other YeiH-specific CD8+ T cells regardless of whether they emerge in patients or healthy individuals.

To determine the transcriptional differences that make the pathogenic-containing effector cluster (Cluster 4) unique from the non-pathogenic, terminally differentiated effector cluster (Cluster 3), we performed differential gene expression analysis. The pathogenic-containing cluster expressed 74 differentially expressed genes (DEGs), including *KLRB1* (encoding CD161), *SLC4A10, LTB, CEBPD, CCR6*, and *IL7R*, while the non-pathogenic effector cluster showed higher expression of *CCL5* (**Fig. 3F**), consistent with our prior transcriptional analysis of pathogenic CD8+ T cells(*9*). We therefore curated a “consensus” set of twenty genes (after excluding the TCR genes *TRAV21* and *TRBV9*) that distinguish pathogenic CD8+ T cells from other cells that bind the YeiH-antigen (current data) or from other CD8+ T cells that also express CD161 (prior data)(*9*) **(Fig. 3G).** These 20 genes are hereafter referred to as a “pathogenic gene signature” for CD8+ T cells in AAU and axSpA.

We next asked whether the pathogenic gene signature bore transcriptional similarities to previously characterized T cell subtypes. The pathogenic signature bore significant similarity to a list of marker genes that characterized CD161^int^CD8^+^ T cells(*29*); 12 of the 20 pathogenic signature genes were present in the CD161^int^CD8^+^ T cell marker gene list (p < 0.01) (**Data File S3**). Accordingly, the CD161^int^CD8^+^ T cell signature had the highest average activity in the pathogenic cluster (**Fig. 3H**) (*30*). This analysis confirms that the transcriptional landscape of the pathogenic cells in the peripheral blood of patients with axSpA and AAU bears similarities to previously described gut-resident CD161^int^CD8^+^ memory T cells.

### Transcriptional overlap of pathogenic cells across tissues in axSpA, AAU, and PsA

As the pathogenic gene signature was identified via sorted CD8+ T cells from the blood, we investigated whether this signature was preserved in CD8+ T cells entering inflamed tissues (joint, eye) of patients with HLA-B*27-associated diseases. We chose to focus on axSpA, AAU, and PsA, another disease known to be associated with HLA-B*27(*1*). We first tested whether pathogenic TCRs could be detected in PsA. We re-analyzed a bulk TCRꞵ sequencing dataset of CD8+ T cells from the blood and synovial fluid of axSpA and PsA patients(*8*). Given the lack of TCRα sequencing information in the dataset, we used the CDR3ꞵ motif and *TRBV9/5-5/5-4* (Criteria 2) to define pathogenic TCRs. We found pathogenic TCRs in both the blood and synovial fluid of axSpA and PsA patients, with higher frequencies in the synovial fluid than in the blood (**Fig. 4A**). The detection of the broadened pathogenic TCRꞵ motif in PsA patients suggests an underlying shared biology across HLA-B*27-associated diseases.

**Fig. 4.**
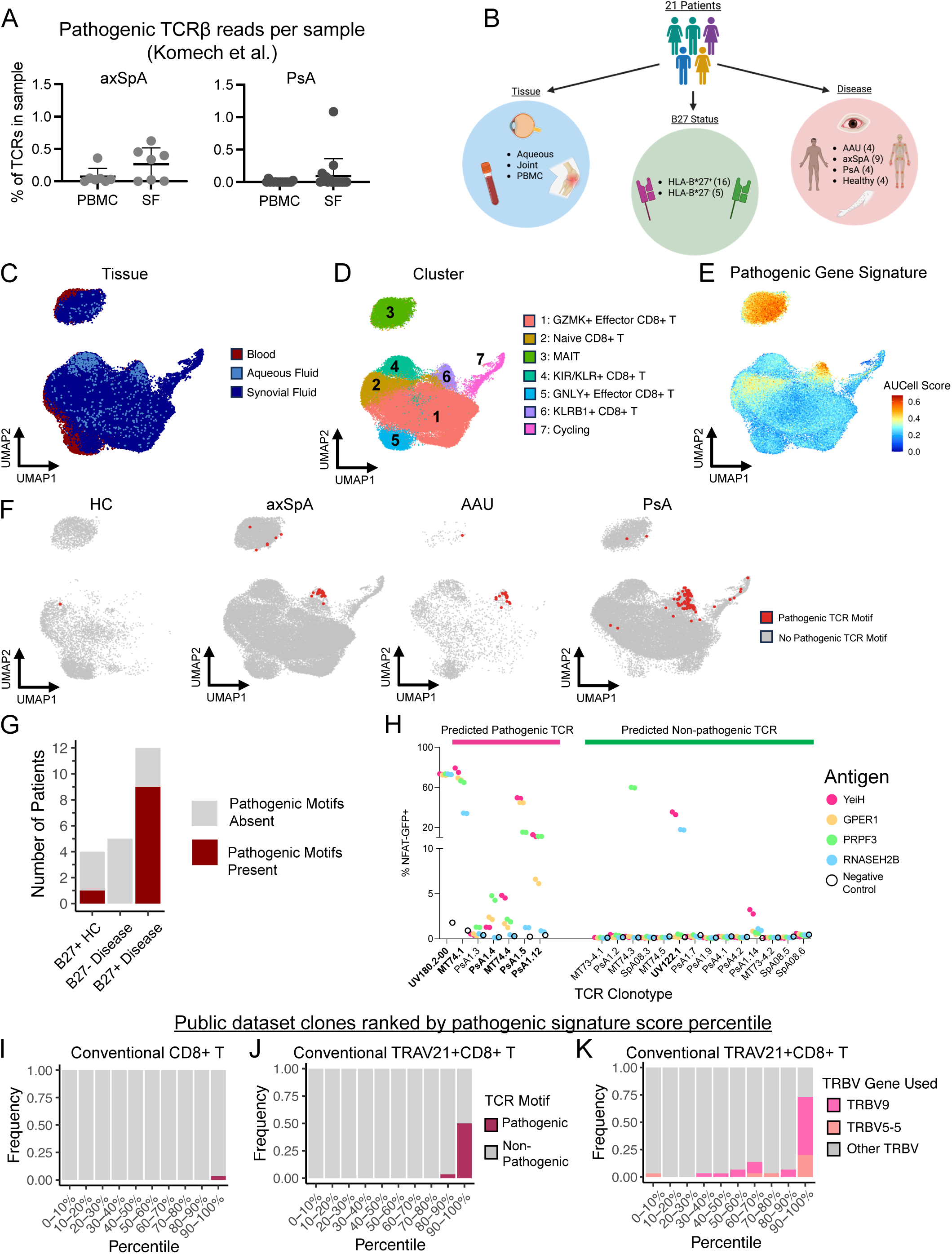
Pathogenic gene signature identifies pathogenic TCRs in public datasets of axSpA/AAU. **(A)** Percentage of reads containing the pathogenic TCR motif in peripheral blood (PBMC) and synovial fluid (SF) from patients with axSpA and PsA. Error bars indicate mean and standard deviation. **(B)** Illustration summarizing the four public CD8+ T cell scRNA-seq + scTCR-seq datasets re-analyzed in panels (C) to (K). **(C** to **E)** UMAP plots of the combined scRNA-seq data from the 4 public datasets. Plots are colored by tissue (C), cell type (D), and expression of the pathogenic gene signature from Fig. 3 (E). **(F)** UMAP plots, split by disease, displaying the location of CD8+ T cells with the pathogenic TCR motif (red). **(G)** Barplot of the number of donors containing pathogenic TCR motifs in the 4 public single-cell datasets split by HLA-B*27 and disease status. **(H)** NFAT-GFP reporter activity for selected Cluster 6 TCRs after exposure to antigen-loaded HLA-B*27+ K562 cells. UV180.2 is used as a positive control. Cross-reactive TCRs are shown in bold on the X-axis. TCRs were considered cross-reactive if they reacted to YeiH and one or more human autoantigens (GPER1, PRPF3, RNASEH2B). **(I** to **K)** Clonotypes from the 4 public datasets, ranked in order of their average pathogenic gene signature score (E) and binned into corresponding percentiles. Clonotypes with pathogenic TCR features are highlighted. Barplots show the pathogenic TCR motif (maroon) in conventional CD8+ T cells (I); the pathogenic TCR motif in conventional TRAV21+CD8+ T cells (J); and *TRBV9* (pink) or *TRBV5-5* (orange) in conventional TRAV21+CD8+ T cells (K).

Next, we compiled four published scRNA-seq datasets of 79,777 CD8+ T cells from 17 patients and 4 healthy controls (*9, 19, 31, 32*). The cells in this public dataset were isolated from peripheral blood (HC, axSpA, PsA, AAU), synovial fluid (axSpA, PsA), and aqueous fluid (AAU) from 16 HLA-B*27+ individuals (from axSpA, AAU, PsA, and healthy) and 5 HLA-B*27-individuals (from axSpA and PsA) (**Fig. 4C, Fig. S4B, Data File S4**). We found 10 clusters corresponding to 7 distinct CD8+ T cell subtypes, including *TRAV1-2*+ mucosal-associated invariant T (MAIT) cells (cluster 3) and conventional *KLRB1*+ CD8+ T cells (cluster 6) (**Fig. 4D and Fig. S4, C and D**). Conventional *KLRB1+* cells made up less than 1% of the peripheral blood cells in both healthy controls and patients, but were expanded in the synovial and aqueous fluid (**Fig. S4E**). We found that the pathogenic signature displayed the greatest expression in two specific clusters: MAIT cells (cluster 3) and conventional *KLRB1*+ CD8+ T cells (cluster 6) (**Fig. 4E**). Analysis of the paired scTCR-seq data confirmed the presence of MAIT cell TCR sequences in cluster 3 and revealed that the cells with pathogenic TCR sequences localized mainly to cluster 6 (**Fig. 4F, Fig. S4F).** The pathogenic cells clustered together regardless of dataset, tissue of origin, or disease, suggesting that their gene expression profile is distinct, consistent, and reproducible across various contexts (**Fig. S4, G and H**). Importantly, pathogenic TCRs were present only in HLA-B*27+ individuals **(Fig. 4G).**

To provide proof of concept that our *in silico* TCR pathogenicity predictions were accurate in the public datasets, we selected 6 predicted pathogenic *TRAV21+* TCRs and 14 predicted non-pathogenic *TRAV21+* TCRs from cluster 6 to test with the Jurkat NFAT-GFP reporter system (**Data File S5**) (*21*). We chose TCRs from across the range of pathogenic gene score values present in Cluster 6. Five of six predicted pathogenic TCRs reacted to both YeiH and at least one autoantigen, while 11/14 predicted non-pathogenic TCRs did not respond to either (**Fig 4H**). Among the predicted non-pathogenic TCRs, MT74.3 reacted to PRPF3 but not YeiH. By contrast, PsA1.14 displayed a modest response to YeiH. UV122.1, the only predicted non-pathogenic TCR with a robust reaction to both YeiH and an autoantigen (RNASEH2B), displayed a *TRAV21-*containing alpha chain paired to a beta chain containing *TRBV7-3* (not tested in our screen) and the CDR3ꞵ motif “CASSPG**G**YSTDTQYF”, which differs from our CDR3ꞵ search criteria only at position 7. Our predicted pathogenic TCR validation provides evidence that the pathogenic TCR criteria performs well in single-cell TCR analysis.

### MAIT and pathogenic cells express similar gene programs

As MAIT cells also express the pathogenic signature, we investigated which genes were shared. We performed differential gene expression analysis on the MAIT cell population and compared the resulting DEGs to the pathogenic signature (**Fig. S4I**). Eight genes were shared between the MAIT cell DEGs and the pathogenic gene signature: *TNFRSF25, IL7R, AQP3, LTB, CEBPD, KLRB1, JAML,* and *RUNX2* (**Fig. S4J**). This overlap was significant (padj < 0.001) according to Fisher’s Exact Test (**Data File S3**). The overlapping gene expression of pathogenic CD8+ T cells and MAIT cells is consistent with prior reports of transcriptional overlap between CD161^int^ conventional αꞵT cells and CD161^hi^ MAIT cells(*29*).

### Pathogenic gene signature identifies CD8+ T cells with cross-reactive TCRs

We next assessed whether the pathogenic gene signature could enrich for CD8+ T cells with a pathogenic TCR. For each TCR clonotype in the public datasets, we calculated the average pathogenic gene signature score across all the cells belonging to the TCR (**Fig. 4E)**. We removed the MAIT cell cluster from our calculations to focus on conventional CD8+ T cells. We then ranked the remaining clonotypes in the dataset in order of their pathogenic signature expression and assigned them each a percentile. All clonotypes containing a pathogenic CDR3ꞵ motif scored above the 90^th^ percentile for pathogenic signature expression, indicating a correlation between pathogenic TCR features and the pathogenic gene module (**Fig. 4I**).

Given that the *TRAV21* gene segment is a consistent feature of pathogenic TCRs(*7, 9*), we also performed a sub-analysis of only *TRAV21*+ clonotypes, re-calculating the percentiles using only the *TRAV21+* conventional CD8+ T cells. Amongst the clonotypes that were 90^th^ percentile or above for pathogenic signature expression in this analysis, ∼40% displayed a pathogenic TCR. Meanwhile, no clonotypes below the 80^th^ percentile displayed the pathogenic CDR3ꞵ motif (**Fig. 4J**). High-ranking *TRAV21+* clonotypes also disproportionately used *TRBV9* or *TRBV5-5* (**Fig. 4K**). This TRBV gene usage pattern was not observed in the analysis of all conventional CD8+ T cells, suggesting that *TRBV9* and *TRBV5-5* are only associated with pathogenic activity when paired with *TRAV21* (**Fig. S4K**). Furthermore, a randomly selected gene set of 20 genes showed no ability to stratify clonotypes by pathogenic TCR features or TRBV gene usage (**Fig. S4, L to O**). The specific correlation of the pathogenic gene signature with pathogenic TCR features suggests a fundamental connection between the TCR specificity of the pathogenic cells and their specific gene expression program.

### Pathogenic CD8+ T cells can be detected in the gut of HLA-B*27+ individuals

Prior work has raised the possibility that HLA-B*27-associated pathogenic CD8+ T cells may undergo their first antigen-specific response in the gut. However, detection of pathogenic CD8+ T cells in the intestine has not yet been reported. To determine whether pathogenic cells can be found in the gut, we initiated a campaign to collect intestinal biopsies from HLA-B27+ individuals. We focused our recruitment on HLA-B*27+ patients with existing or suspected inflammatory bowel disease (IBD) due to the increased frequency of clinically indicated colonoscopies with biopsy.

We obtained samples of peripheral blood and gut (ileum or colon) tissue at the time of colonoscopy (**Fig. 5A, Data File S6**). We screened the Washington University electronic medical record (44,078 patients) for the co-occurrence of HLA-B*27 and IBD; in parallel, we also screened 992 patients from the Washington University Digestive Diseases Research Cores Center (DDRCC) for the presence of the HLA-B*27 allele via PCR testing. We identified 113 HLA-B27+ individuals across the two screening strategies. After removing duplicates, deceased patients, and those meeting exclusion criteria (see Methods), 99 candidates remained. Of these HLA-B*27+ IBD individuals, we collected three paired samples of intestinal tissue and peripheral blood. In addition, we added one patient with HLA-B*27+ AAU who was enrolled at a diagnostic colonoscopy but ended up showing no evidence of IBD. We sorted CD8+ T cells from the intestinal biopsies from each sample for paired scRNA-seq and scTCR-seq. In parallel, we sorted YeiH+CD8+ T cells from the 4 paired blood samples to enrich for pathogenic CD8+ T cells. We also added 2,482 bulk CD8+ T cells from 2 of the 4 blood samples to the ∼2,220 YeiH-sorted CD8+ T cells. After scRNA-seq, we recovered 3,148 cells (1,192 blood, 1,956 gut). We identified 8 clusters in the data, corresponding to 5 discrete cell types (**Fig. 5B, Fig. S5, A and B**). Naïve/memory CD8+ T cells, GZMK+ effector CD8+ T cells, and conventional KLRB1+ CD8+ T cells could be found in both the peripheral blood and the gut, while KIR+ CD8+ T cells were predominantly gut-derived, and GNLY+ CD8+ T cells were predominantly blood-derived (**Fig. 5C, Fig. S5C**). TCR analysis identified pathogenic cells in the gut tissue of all three IBD participants, with two of the three also displaying pathogenic TCRs in the peripheral blood (**Fig. S5D**). We also detected pathogenic TCRs in the peripheral blood of the AAU non-IBD participant, although the number of pathogenic TCRs in the gut were below the level of detection. Regardless of tissue, pathogenic cells localized almost exclusively (98%) to the KLRB1+ cluster (**Fig. 5D**). Concordant with our YeiH-sorted dataset and our aggregated public dataset, the KLRB1+ cluster had the highest pathogenic signature expression (**Fig. 5, E and F**). Furthermore, within the KLRB1+ cluster, the IBD patients displayed significantly (p < 0.05) higher average pathogenic signature expression in the gut than in the peripheral blood, while the one patient without IBD did not display this pattern (**Fig. 5G**).

**Fig. 5.**
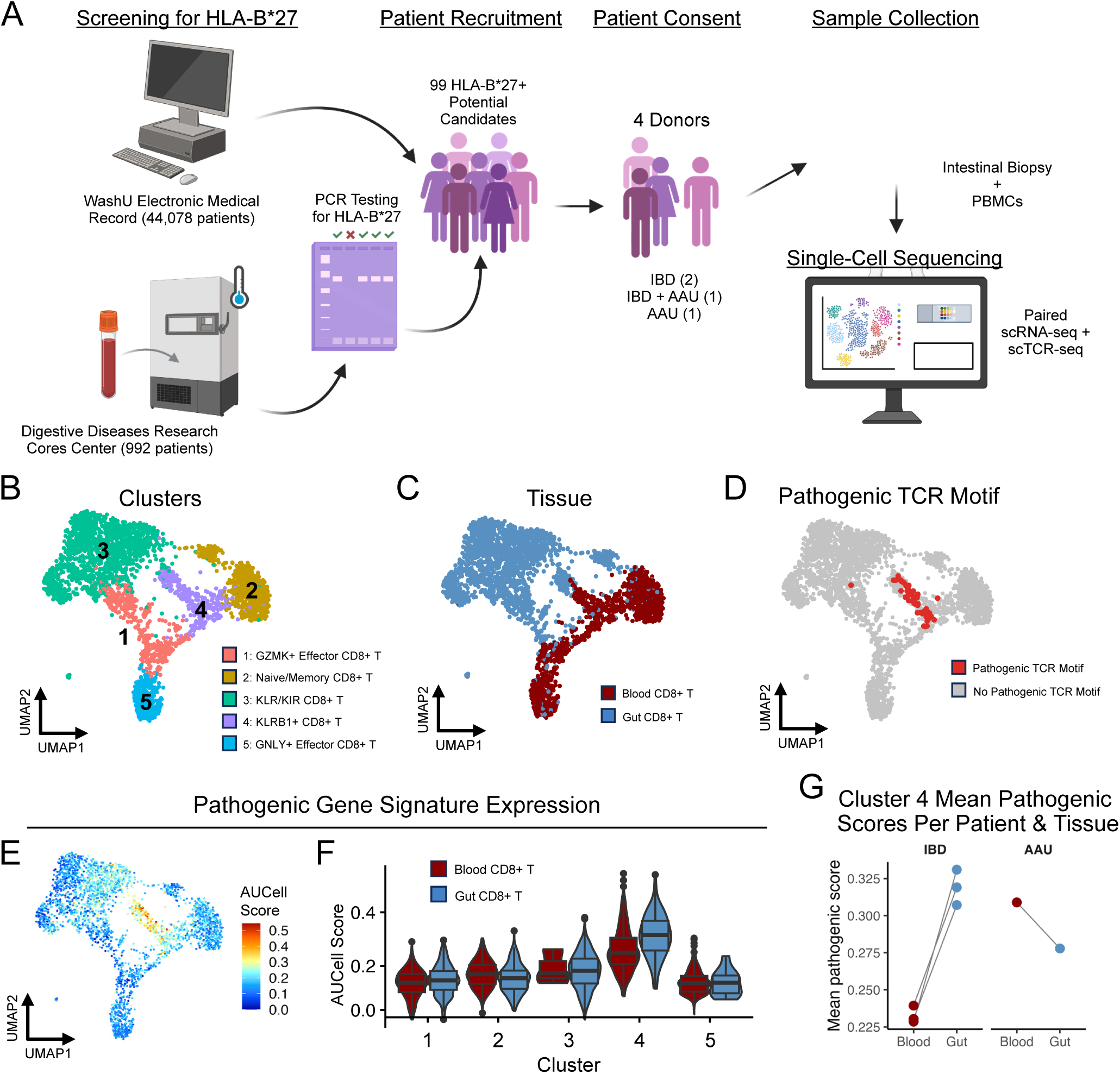
Pathogenic CD8+ T cells are present in the gut of HLA-B*27 individuals. **(A)** Schematic depicting sample collection for the paired gut and blood CD8+ T cell sort. **(B** to **E)** UMAP plots of the paired scRNA-seq + scTCR-seq analysis of gut and blood CD8+ T cells from HLA-B*27+ individuals (3,148 cells total). Plots show cluster information (B), sample location (C), presence of the pathogenic TCR motif (D), and AUCell activity scores for the pathogenic gene signature (E). **(F)** Violinplots of the pathogenic gene signature expression shown in panel (E). Violinplots are split by cluster and sample location. **(G)** Mean per-sample pathogenic signature expression score within Cluster 4 (the KLRB1+ cluster). Samples are split by location (blood, red; gut, light blue).

Collectively, these data demonstrate that pathogenic TCRs are found within the gastrointestinal tract and suggest that these cells may acquire their specific transcriptional program at this initial site of antigen activation.

### Pathogenic CD8+ T cells appear to produce type 17 cytokines *in vitro*

We and others have previously reported that pathogenic CD8+ T cells express CD161 (*KLRB1*), which is a marker of type 17 immune cells(*9, 19, 20, 29*). Therefore, we sought to determine whether the pathogenic CD8+ T cells, which may receive programming in the gut, also expressed an overall “type 17-like” phenotype.

We searched for canonical type 17 genes in our analysis of the YeiH+CD8+ T cell blood dataset (**Fig. 3**), the public joint/eye/blood dataset (**Fig. 4**), and the HLA-B*27+ IBD gut/blood dataset (**Fig. 5**). Compared to the other conventional CD8+ T cells, the KLRB1+ (pathogenic TCR motif-containing) clusters displayed increased expression of *RORC* and *IL23R*, both central genes in type 17 differentiation(*33, 34*) (**Fig. 6, A to C**). *IL17A* was detected at low levels in the public joint/eye/blood dataset and the HLA-B*27+ IBD gut/blood dataset mainly in the pathogenic TCR motif-containing clusters (**Fig. 6, A to C**). *IL-22* was detected in the HLA-B*27+ IBD gut, consistent with prior reports of increased IL-22 expression in the gut(*35, 36*) (**Fig. S6, A to B**). All pathogenic TCR motif-containing clusters also displayed increased activity of a Th17 signature(*37*) (**Fig. 6, D to F**).

**Fig. 6.**
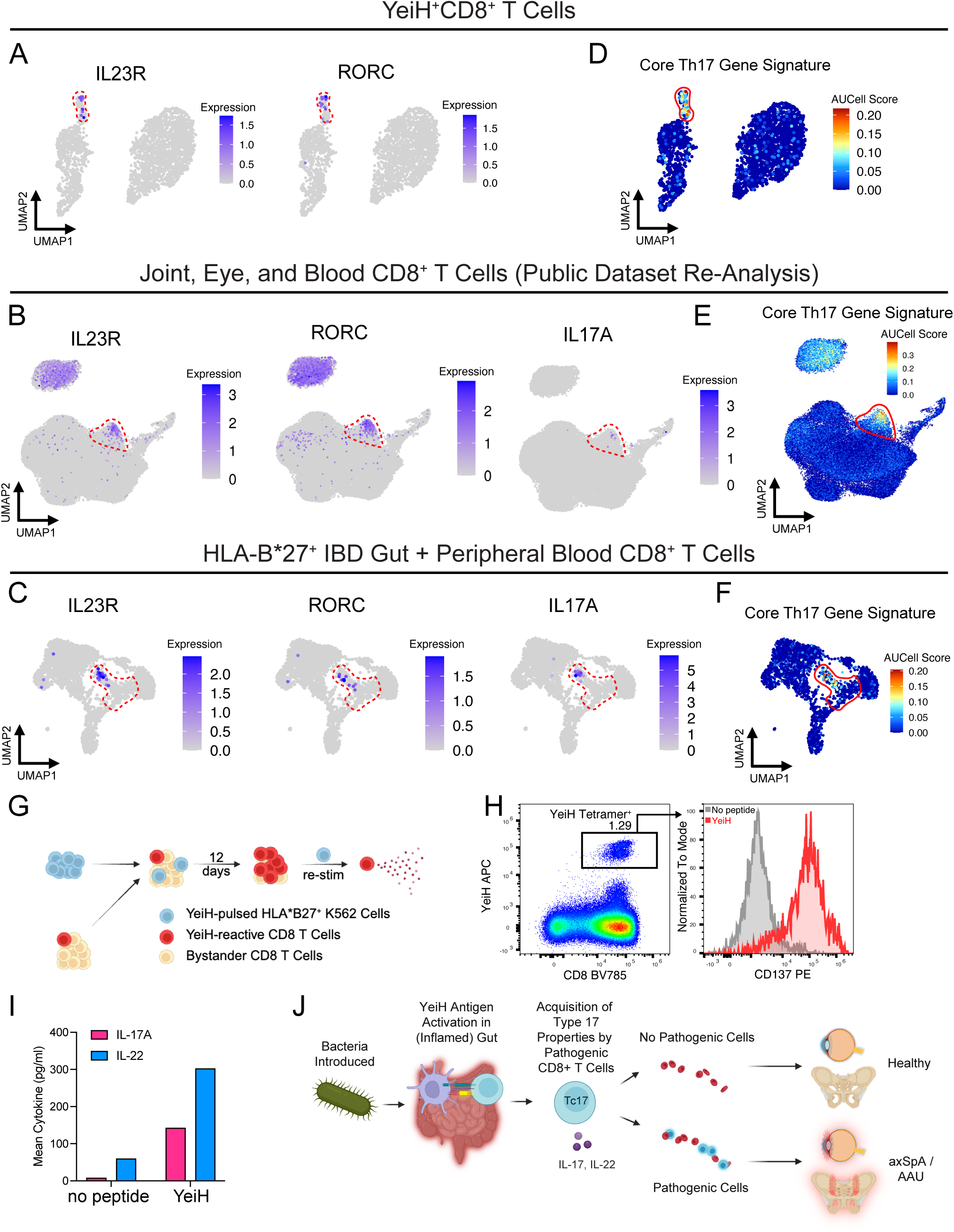
Pathogenic CD8+ T cells express Type 17 genes and produce Type 17 cytokines. **(A)** UMAP plots depicting expression of *IL23R* and *RORC* in the YeiH+CD8+ T cells from Fig. 3. KLRB1+ cluster is indicated (red dashed line). **(B** and **C)** UMAP plots depicting expression of *IL23R, RORC,* and *IL17A* in (B) the joint, eye, and blood CD8+ T cells from Fig. 4 and (C) the HLA-B*27+ IBD gut and peripheral blood CD8+ T cells from Fig. 5. KLRB1+ clusters are indicated (red dashed lines). **(D** to **F)** UMAP plots depicting AUCell activity score for the “Core Th17” gene signature from Ramesh et al. Datasets shown are the YeiH+CD8+ T cells (D), the joint, eye, and blood CD8+ T cells (E), and the HLA-B*27+ IBD gut and peripheral blood CD8+ T cells (F). KLRB1+ clusters are indicated (red lines). **(G)** Schematic of the YeiH+CD8+ T cell cytokine panel. **(H)** Representative sorting strategy for the YeiH+CD8+ T cells. **(I)** IL-17A and IL-22 production of YeiH+CD8+ T cells after stimulation with HLA-B*27+ K562 cells alone (no peptide) or pulsed with YeiH.232-240. Data is the mean of two biological replicates. **(J)** Illustration of the proposed model for axSpA and AAU pathogenesis.

To test whether the pathogenic cells may produce type 17 cytokines *in vitro*, we developed a platform to identify the cytokines released by antigen-specific CD8+ T cells. Given that YeiH+ CD8+ T cells make up a low percentage of the total CD8+ T cell population (**Fig. 2A**), we modified a prior assay(*9*) by co-culturing purified CD8+ T cells with YeiH-pulsed HLA-B*27^+^ K562 cells to maximize antigen-specific T cell activation and expansion (**Fig. 6G**). This expansion occurs in the absence of polarizing cytokines. After ∼12 days of T cell proliferation, we re-stimulated CD8+ T cells with YeiH-pulsed HLA-B*27^+^ K562 cells for 16 hours and verified antigen-specific activation of CD8+ T cells via CD137 upregulation (**Fig. 6H**)(*9*). We then tested for the antigen-specific release of effector molecules into the culture supernatant using a multiplex assay for 27 different cytokines and granzymes. IL-17A and IL-22 were the two cytokines with the greatest fold change in the media in response to YeiH stimulation (**Fig. 6I, Fig. S6C**). In contrast, YeiH stimulation led to only a moderate increase in IFN-g, suggesting that pathogenic CD8+ T cells may promote a Type 17 response (**Fig. S6C**). Collectively, these transcriptional and functional data suggest pathogenic CD8 T cells may develop a Type 17 response in the intestinal tract and retain that program after entering the eye and the joint (**Fig 6J**).

## DISCUSSION

Here, we refine our understanding of the TCR sequences and gene expression of pathogenic CD8+ T cells in HLA-B*27+ axSpA and AAU. Moreover, we provide evidence that pathogenic cells may harbor a Type 17 transcriptional program and show that these pathogenic cells can be found in the colon of HLA-B*27+ individuals. Collectively, these data support the model that the pathogenic CD8+ T cells first encounter antigen in the gut and travel to the eye and joint to initiate a pathologic Type 17 response.

The pathogenic cells in axSpA and AAU have been reported to use the TRBV gene segments *TRBV5-5* and *TRBV9* for their TCRs(*7, 9*). Our TRBV mutagenesis experiments suggest that TRBV5-5 and TRBV9 are likely interchangeable for antigen reactivity, while TRBV5-4 also may produce pathogenic TCRs to a lesser extent. Mutagenesis of the CDR3ꞵ sequence suggests that the canonical CDR3ꞵ motif “[L/T]-[Y/F]-S-T” may not encompass all possible pathogenic sequences. Although CDR3ꞵ positions 8-10 ([Y/F]-S-T) appear necessary for antigen cross-reactivity, positions 5 through 7 could accommodate greater amino acid variability while preserving cross-reactivity. Collectively, our data suggest that the pathogenic CD8+ T cells have a broader TCR repertoire than previously appreciated. Identifying the full range of pathogenic TCR sequences has important clinical implications when using high-throughput sequencing methods to stratify patients for T cell depletion therapy by seniprutug. Importantly, if the entire pathogenic cell population is not depleted, surviving pathogenic CD8+ T cell clones may lead to treatment failure.

Our broadened definition of pathogenic TCRs may be important in supporting a clinical diagnostic. Using peripheral blood samples, we provide proof-of-concept that a simple presence/absence cutoff for pathogenic TCRs can distinguish patients from healthy controls with over 80% sensitivity and specificity. These data do contain several limitations. (1) The number of participants tested were small, so the exact sensitivity and specificity metrics may change in larger cohorts. (2) TCR-based tests work poorly if a patient’s circulating pathogenic population does not exceed the limit of detection. However, as high-throughput TCR sequencing strategies become more economical and robust, and as our understanding of the pathogenic TCR repertoire becomes more complete, it is likely that a TCR-based test like TIRTL-seq could become a useful clinical tool. Possible uses include screening (in the general practitioner’s office), diagnostics (in the rheumatologist’s office), or as a stratifying tool to identify patients who might be eligible for T cell depletion therapy.

The analysis of paired scTCR-seq and scRNA-seq data from our own in-house dataset (axSpA and AAU) as well as four publicly available datasets (axSpA, AAU, and PsA) revealed that cells with pathogenic TCRs form a distinct cluster and display a unique transcriptional signature. Despite the lack of TCR genes present in the pathogenic gene signature, conventional CD8+ T cells with a high pathogenic gene signature score were more likely to bear the pathogenic TCR motif than their low-scoring counterparts, providing a clear connection between TCR specificity and transcriptional programming. This transcriptional signature is reproducible across datasets, tissues, and HLA-B*27-related diseases, suggesting that HLA-B*27-restricted antigens may be a singular driving mechanism across multiple types of arthritis and AAU. While current disease classification systems combine patients regardless of HLA-B*27 status, a future HLA-B*27-based re-classification may become important for clinical treatment guidelines based on disease pathogenesis.

The current model posits that the pathogenic cells are initially programmed in the gut as a response to Gram-negative bacteria such as *E. coli, Salmonella,* and *Shigella* spp(*7*). Prior studies have detected subclinical levels of gut inflammation in axSpA patients, suggesting that enteric inflammation may play a role in the activation or programming of pathogenic CD8+ T cells(*12, 13*). We detected pathogenic TCRs in gut biopsies from all three of the HLA-B*27+ Crohn’s disease patients in our cohort, however, pathogenic TCRs were below the level of detection in the patient with AAU. In parallel, the pathogenic gene signature was increased in the gut over the peripheral blood in the patients with Crohn’s disease, but not in the HLA-B*27+ / AAU+ / IBD-patient, suggesting that the pathogenic signature may be augmented by intestinal inflammation. Thus, intestinal inflammation may promote the development of axSpA and AAU through both the programming and expansion of pathogenic CD8+ T cells.

How and where pathogenic CD8+ T cells become programmed remains an open question. Previous work has found shared CD8+ T cell clones in the blood, skin, and joint of PsA patients, suggesting that CD8+ T cells can travel between tissues to promote inflammation in multiple locations (*38*). The pathogenic cells express the same gene signature in the gut as they do in the peripheral tissues, suggesting that their distinct transcriptional profile originates in the gut (as opposed to being acquired somewhere between the gut and the joint or eye). However, we cannot exclude the possibility of pathogenic CD8+ T cells returning to the gut after exposure to autoantigens in other tissues.

The pathogenic cell transcriptome bears similarities to three previously defined, interconnected cell types: Th17 cells, MAIT cells, and CD161^int^ CD8+ T cells(*20, 29, 37*). MAIT cells and CD161^int^ CD8+ T cells have both been shown to display mixed Type 17 and Type 1 effector capabilities(*29, 39*). Accordingly, we find that the pathogenic CD8+ T cells express key Type 17 marker genes and are enriched for a Th17 signature in single-cell analysis. We also find that YeiH peptide stimulation of patient-derived CD8+ T cells elicits moderate amounts of IFN-g and high amounts of IL-17A and IL-22 *in vitro*. The Type 17 capabilities of the pathogenic cells suggest that their role may involve coordinating the inflammatory environment in HLA-B*27-associated diseases. IL-17 inhibitors have already been shown to be effective for treating axSpA in the clinic, and targeting the pathogenic cells can induce remission in a proof-of-concept case study, suggesting that the cells are at the center of the inflammatory milieu and that their removal results in the de-escalation of the inflammatory environment(*11, 15*). In this model, HLA-B*27 fosters the development of cross-reactive, Type 17 pathogenic cells, which carry a Type 17 inflammatory response from the gut to the eye or the joint, likely recruiting other Type 17 cells like Th17s. This model may explain why patients with HLA-B*27^+^ axSpA might respond to both IL-17 inhibitors and antibody-targeted depletion of pathogenic TCRs. Our expanded understanding of the TCRs and function of pathogenic CD8+ T cells reveals the shared biology underlying HLA-B*27-associated diseases, paving the way for novel diagnostics that would support early disease detection and treatment.

## MATERIALS AND METHODS

### Study Design

This study aimed to characterize the TCR repertoires, transcriptomes, and functional characteristics of pathogenic CD8+ T cells in HLA-B*27-associated axSpA and AAU. We collected CD8+ T cells from peripheral blood and intestine of HLA-B*27+ healthy individuals and HLA-B*27+ participants with axSpA, AAU, and IBD. Participants with HLA-B*27+ AAU fulfilled the Standardization of Uveitis Nomenclature Working Group classification criteria for HLA-B*27+ AAU(*40*). Participants with HLA-B27+ AS satisfied the modified New York criteria(*41*). Participants with HLA-B*27+ axSpA satisfied the 2009 SpondyloArthritis International Society (ASAS) classification criteria for axial spondyloarthritis(*42*). Participants with IBD were included based on a clinical diagnosis of Crohn’s disease. Healthy controls were recruited through the Volunteers for Health registry at Washington University in St. Louis. Healthy controls were free of autoimmune diseases. HLA-B*27 positivity was confirmed through PCR in the laboratory or clinically indicated HLA testing.

We performed bulk and single-cell RNA-seq and TCR-seq and cytokine production panels on the CD8+ T cells collected from the participants. We also performed saturation mutagenesis experiments on cells from the J76 Jurkat cell line. For the saturation mutagenesis experiments, two technical replicates were produced per library. For each of the high-throughput TCR-seq datasets, at least four biological replicates were produced per condition (see Results). For the single-cell sequencing data, at least 3 biological replicates were collected in each tissue for each dataset; per-dataset numbers are detailed in the associated figures. For the Luminex cytokine panel, two biological replicates were produced per condition.

### Study Approval

All human participants were enrolled in accordance with Declaration of Helsinki principles and using protocols approved by the institutional review board of Washington University in St. Louis (IRB 201111078, 201704141, 201912043, 202011161, and 202212047).

### Statistical Analysis

Statistical analyses were performed as described using R v4.4.0 and GraphPad Prism v11.0.2. Quantitative data are presented as mean ± SD. For mutational enrichment analysis, Student’s t-test was used to determine statistical significance. For comparisons between two groups (healthy vs. autoimmune), a Wilcoxon Rank-Sum (Mann-Whitney U) test was used. For DEG analysis, the MAST or DESeq2 algorithm was used. Asterisk key for figures: * indicates P-value ≤ 0.05, ** indicates P-value ≤ 0.01.

## List of Supplementary Materials

Materials and Methods

Fig S1 to S8

Data files S1 to S10

References (*40–58*)

## Acknowledgments

We thank GTAC@MGI and GESC@MGI (Washington University School of Medicine) for their help producing cell lines and sequencing data. We thank Dr. Jennifer Foltz and Dr. Wayne Yokoyama for advice, manuscript review, and scientific discussion. We also thank Dr. Nathan Singh for the J76 Jurkat cell line. The schematic cartoons in Figures 1-6 were created with BioRender.com.

## Funding

National Institutes of Health grant F30AI200188 (IR)

National Institutes of Health grant T32HG000045 (IR)

Goldberg Family Foundation (MG, OLG)

National Institutes of Health grant R01CA301054 (MG, OLG)

National Institutes of Health grant K08AR079593 (MP)

National Institutes of Health grant P30AR073752 (MP)

Department of Defense grant AT240082 (MP)

Spondylitis Association of America Rheumatology Research Foundation

Edward Mallinckrodt, Jr. Foundation Arthritis National Research Foundation Doris Duke Foundation

Intestinal Biopsy collection was supported by the Biobank and Big Data Core within the Digestive Diseases Research Cores Center (under National Institutes of Health grant P30DK052574)

## Author contributions

I.R., M.G., O.L.G., and M.A.P. designed the study, analyzed the data, and wrote the manuscript. X.Y. contributed to the design of the TCR mutational analysis. Q.W. performed TCR mutational analysis. Q.W., R.D., S.Y.L., P.H.A.S., and A.B. performed HLA-B*27 screening of participants and collected blood and intestinal specimens. T.A.P.N., S.W., and P.A.M. performed TIRTL-seq experiments. N.C.B. performed computational analysis of TIRTL-seq. Q.W., R.D., and S.Y.L performed the sorting experiments for scRNA-seq. S.Y.L. performed CD8+ T cell cultures for functional analysis. I.R. performed TCR cloning and lentivirus generation. L.D., M.T., Z.Q., R.I., and L.S.T. collected clinical information for scRNA-seq datasets. All authors reviewed and edited the manuscript.

## Competing interests

M.A.P. has received consulting fees from UCB, Priovant, and EpanaBio, received honorarium from Incyte, received research funding from Merck paid to the university, and holds equity in EpanaBio. M.G. and O.L.G. receive consulting fees from the Jaime Leandro Foundation for Therapeutic Cancer Vaccines and Pathfinder Oncology Inc., and received research funding from Natera Inc. paid to the university. N.C.B. is a scientific advisor to Epana Bio, Inc, a consultant to Columbus Instruments. L.D. is currently an employee of AstraZeneca. L.S.T. has received consultancy or speaker fees and/or research funding from AbbVie, CESAS Medical, GSK, Sanofi A/S, and UCB. The remaining authors declare no other competing interests.

## Data and materials availability

Detailed methods to reproduce the findings in this paper are provided in the Supplementary Materials and Methods and in the original analysis code, which is publicly available on GitHub (https://github.com/Paley-Lab/risch_paley_axSpA_AAU_pathogenic_cd8t/). Supplementary figures and tables are available in Figs. S1-S8 and Data Files S1-S9. Source data for the Figures and Supplementary Figures is available in Data File S10. New sequencing datasets presented in this paper are publicly available as follows: Unprocessed files for the mutational screen, the TIRTL-seq, and the scRNA-seq and scTCR-seq datasets will be uploaded to dbGAP (study number pending). Processed data for the mutational screen and TIRTL-seq, as well as the count matrices and metadata for the scRNA-seq and scTCR-seq datasets, will be made publicly available on Zenodo (doi:10.5281/zenodo.21998424) upon publication. All reasonable requests for materials will be fulfilled by the lead contact, M.A.P.

## Notes

### Competing Interest Statement

The authors have declared no competing interest.

