## Supplemental Figures for "Refinement of the Pathogenic T Cell Receptor Motif Suggests Type 17 CD8+ T Cells Initiate HLA-B*27-associated Autoimmunity"

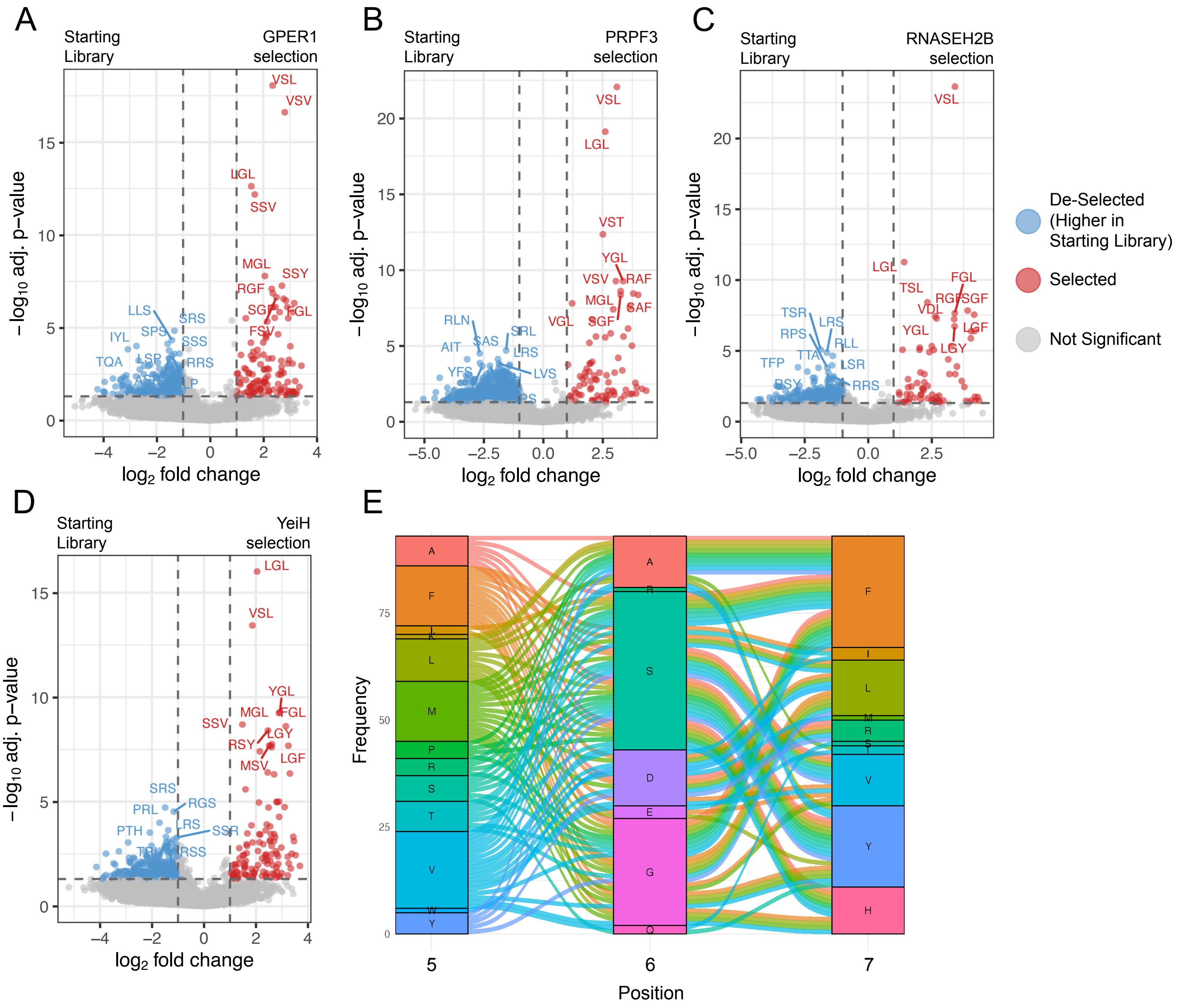
A
Starting Library
GPER1 selection
B
Starting Library
PRPF3 selection
C
Starting Library
RNASEH2B selection
−log10 adj. p−value
log2 fold change
De-Selected (Higher in Starting Library)
Selected
Not Significant
D
Starting Library
YeiH selection
E
Frequency
Position
VSL
VSV
LGL
SSV
MGL
SSY
RGF
SGF
FGL
FSV
LLS
SRS
IYL
SPS
SSS
TQA
LSP
RRS
VST
YGL
RAF
SAF
VGL
RLN
SAS
SRL
AIT
LRS
YFS
LVS
TSL
FGL
TSR
LRS
RPS
RLL
TTA
LSR
TFP
RRS
VDL
LGY
LGF
RGF
SRS
PRL
RGS
PTH
SSR
RSS
SSV
RSY
MSV

**Figure S1.** Quality control and analysis of saturation mutagenesis experiments from Fig. 1. **(A** to **D)** Volcano plots displaying differentially abundant motifs pre- and post- antigen selection in the CDR3β position 5-7 mutational experiment, as per Fig. 1B. Antigens used for selection are GPER1 (A), PRPF3 (B), RNASEH2B (C), and YeiH (D). Cutoffs shown are log2FC > 1 and adjusted p-value < 0.01. Positively selected motifs are shown in red; negatively selected motifs are shown in blue. **(E)** Alluvial plot displaying linkage of amino acids at each position in the cross-reactive motifs shown in Fig. 1G-H.

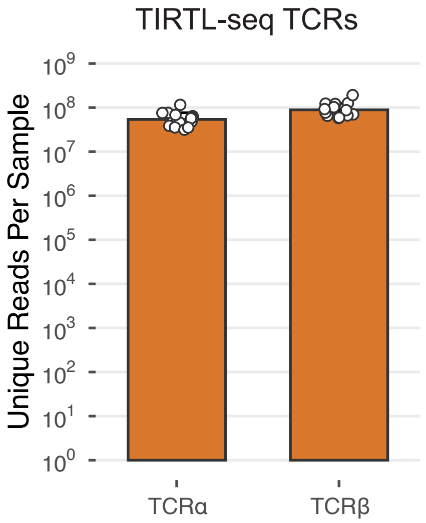


**Figure S2.** UMI counts per sample in TIRTL-seq. Counts for TCRα and TCRβ chains are shown separately. Counts are plotted on a log10 scale. Means plotted are geometric means; error bars are geometric standard deviation. n = 14 TIRTL-seq samples.

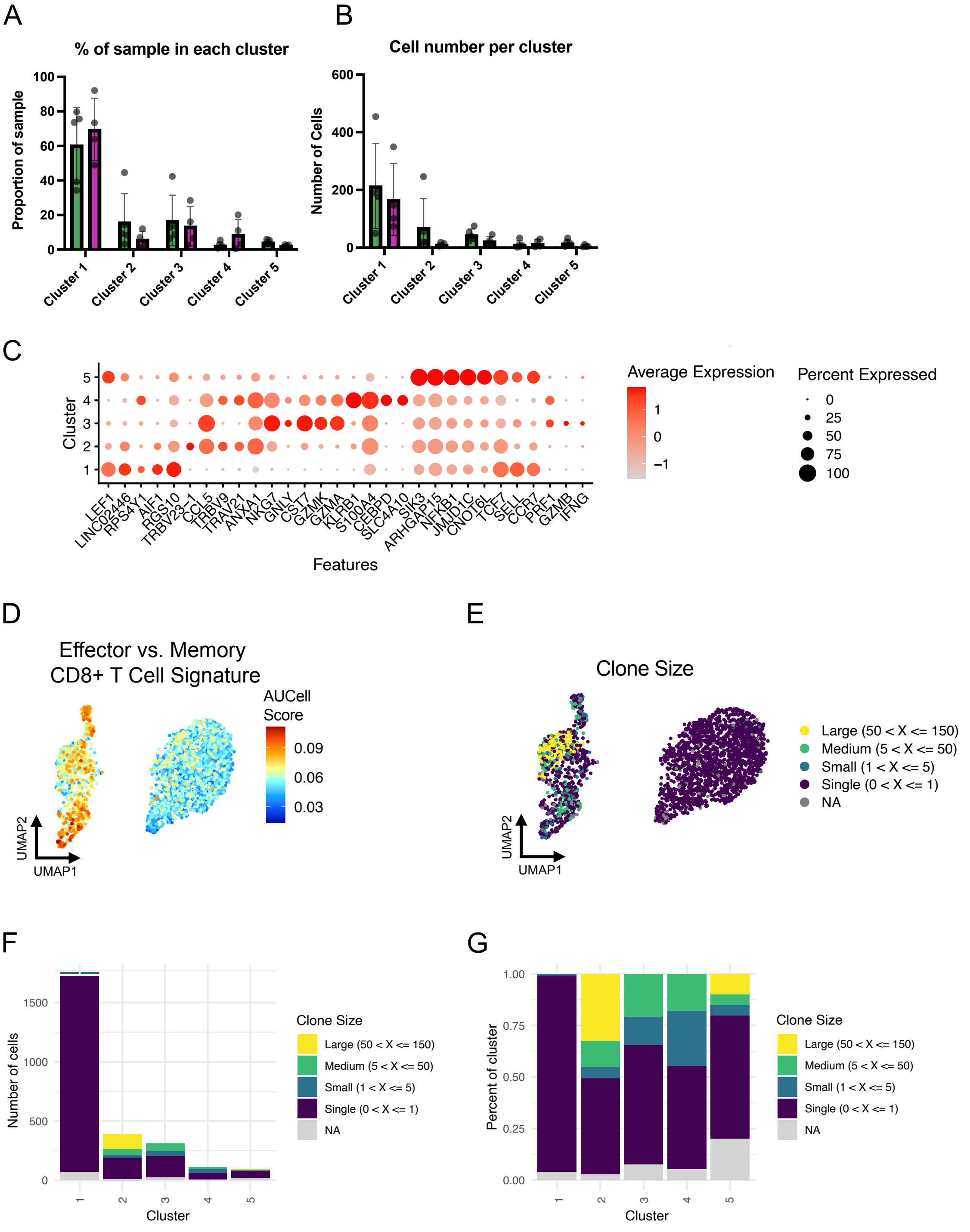

A
% of sample in each cluster
Proportion of sample
Cluster 1
Cluster 2
Cluster 3
Cluster 4
Cluster 5
B
Cell number per cluster
Number of Cells
C
Cluster
Features
LEF1
LINC02446
RPS4Y1
AIF1
RGS10
TRBV23-1
CCL5
TRBV9
TRAV21
ANXA1
NKG7
GNLY
CST7
GZMK
GZMA
KLRB1
S100A4
CEBPD
SLC4A10
SIK3
ARHGAP15
NFKB1
JMJD1C
CNOT6L
TCF7
SELL
CCR7
PRF1
GZMB
IFNG
Average Expression
Percent Expressed
D
Effector vs. Memory
CD8+ T Cell Signature
AUCell Score
UMAP2
UMAP1
E
Clone Size
Large (50 < X <= 150)
Medium (5 < X <= 50)
Small (1 < X <= 5)
Single (0 < X <= 1)
NA
F
Number of cells
Cluster
G
Percent of cluster

**Figure S3.** Analysis of the YeiH+CD8+ T cell scRNA-seq and scTCR-seq data from Fig. 3. **(A)** Cell frequencies (normalized to total cells per sample) for each cluster in Fig. 3B. **(B)** Absolute cell numbers per sample in each cluster in Fig. 3B. **(C)** Dotplot of expression values for selected marker genes, including the top 5 differentially expressed genes (DEGs) per cluster, in the YeiH+CD8+ T cell dataset. DEGs were ranked in order of descending log2FC value; all are significant (padj < 0.01). **(D)** Expression of the MSigDB signature "GOLDRATH_EFF_VS_MEMORY_CD8_TCELL_UP" in the YeiH+CD8+ T cell dataset. Red values indicate higher activity of effector signature genes. Values calculated using the AUCell algorithm. **(E)** UMAP plot of clone size. **(F)** Barplot of absolute cell numbers in each cluster from panel (E). **(G)** Barplot of clone size proportions within each cluster from panel (E).

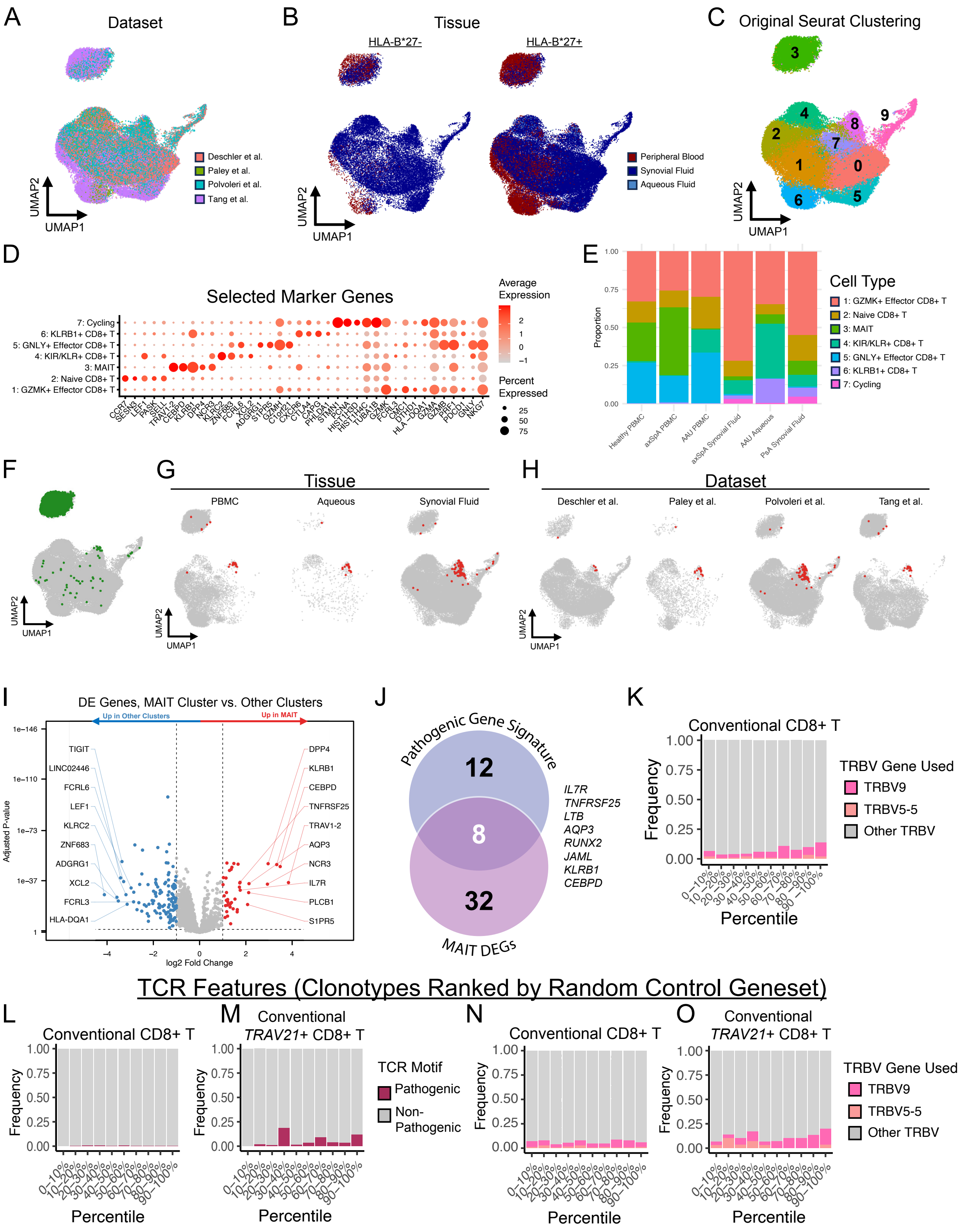
A
Dataset
Deschler et al.
Paley et al.
Polvoleri et al.
Tang et al.
UMAP1
UMAP2
B
Tissue
HLA-B*27-
HLA-B*27+
Peripheral Blood
Synovial Fluid
Aqueous Fluid
UMAP1
UMAP2
C
Original Seurat Clustering
UMAP1
UMAP2
D
Selected Marker Genes
7: Cycling
6: KLRB1+ CD8+ T
5: GNLY+ Effector CD8+ T
4: KIR/KLR+ CD8+ T
3: MAIT
2: Naive CD8+ T
1: GZMK+ Effector CD8+ T
Average Expression
Percent Expressed
E
Proportion
Healthy PBMC
axSpA PBMC
AAU PBMC
axSpA Synovial Fluid
AAU Aqueous
PsA Synovial Fluid
Cell Type
1: GZMK+ Effector CD8+ T
2: Naive CD8+ T
3: MAIT
4: KIR/KLR+ CD8+ T
5: GNLY+ Effector CD8+ T
6: KLRB1+ CD8+ T
7: Cycling
F
UMAP1
UMAP2
G
Tissue
PBMC
Aqueous
Synovial Fluid
UMAP1
UMAP2
H
Dataset
Deschler et al.
Paley et al.
Polvoleri et al.
Tang et al.
UMAP1
UMAP2
I
DE Genes, MAIT Cluster vs. Other Clusters
Up in Other Clusters
Up in MAIT
Adjusted P-value
log2 Fold Change
TIGIT
LINC02446
FCRL6
LEF1
KLRC2
ZNF683
ADGRG1
XCL2
FCRL3
HLA-DQA1
DPP4
KLRB1
CEBPD
TNFRSF25
TRAV1-2
AQP3
NCR3
IL7R
PLCB1
S1PR5
J
Pathogenic Gene Signature
12
8
32
MAIT DEGs
IL7R
TNFRSF25
LTB
AQP3
RUNX2
JAML
KLRB1
CEBPD
K
Conventional CD8+ T
Frequency
Percentile
TRBV Gene Used
TRBV9
TRBV5-5
Other TRBV
TCR Features (Clonotypes Ranked by Random Control Geneset)
L
Conventional CD8+ T
Frequency
Percentile
M
Conventional
TRAV21+ CD8+ T
Frequency
Percentile
TCR Motif
Pathogenic
Non-Pathogenic
N
Conventional CD8+ T
Frequency
Percentile
O
Conventional
TRAV21+ CD8+ T
Frequency
Percentile
TRBV Gene Used
TRBV9
TRBV5-5
Other TRBV

**Figure S4.** Public CD8+ T cell scRNA-seq + scTCR-seq data, re-analyzed from Paley et al., Tang et al., Deschler et al., and Povoleri et al. **(A** to **C)** UMAP plots of the combined datasets, colored by dataset (A), tissue (B), and Seurat clustering before cell type annotation. **(D)** Expression of selected marker genes within each celltype cluster from Fig. 4D. **(E)** Bar plots of the cell type proportions within each disease and tissue. Cell types correspond to Fig. 4D. **(F)** UMAP plot depicting the location of cells with canonical MAIT cell TCR sequences (see Methods). **(G** and **H)** UMAP plots depicting the location of cells with pathogenic TCR motifs. For ease of visualization, UMAPs are split by tissue (G) and dataset (H). **(I)** Volcano plot of the differentially expressed genes (DEGs) in the MAIT cluster (red) versus all other cells in the dataset (blue). All DEGs pass log2FC > 1 and adjusted p-value < 0.01. The top 10 most significant upregulated and downregulated genes according to log2FC are labeled. **(J)** Venn diagram depicting the overlap of the pathogenic gene signature (top) and the MAIT cell upregulated DEGs from panel **I** (bottom). The 8 overlapped genes are shown at right. **(K)** Conventional CD8+ T cell clonotypes from the 4 public datasets were ranked in order of decreasing average pathogenic gene signature score (Fig. 4E) and assigned a corresponding percentile. Barplots represent increasing bins of 10%, with the top percentiles (90%-100%) indicating the clones with the highest average pathogenic gene score. Clones were then highlighted in the barplots according to the presence or absence of the *TRBV9* gene segment (pink) or *TRBV5-5* (orange). **(L** to **O)** Barplots of specified TCR features, as in panel (K); clonotypes in (L) through (O) are ranked according to their expression of a control signature made up of 20 randomly-selected genes. Barplots depict the pathogenic TCR motif (maroon) in conventional clonotypes (L), the pathogenic TCR motif in conventional *TRAV21*+ clonotypes

(M), *TRBV9* and *TRBV5-5* in conventional clonotypes (N), *TRBV9* and *TRBV5-5* in conventional, *TRAV21*+ clonotypes (O).

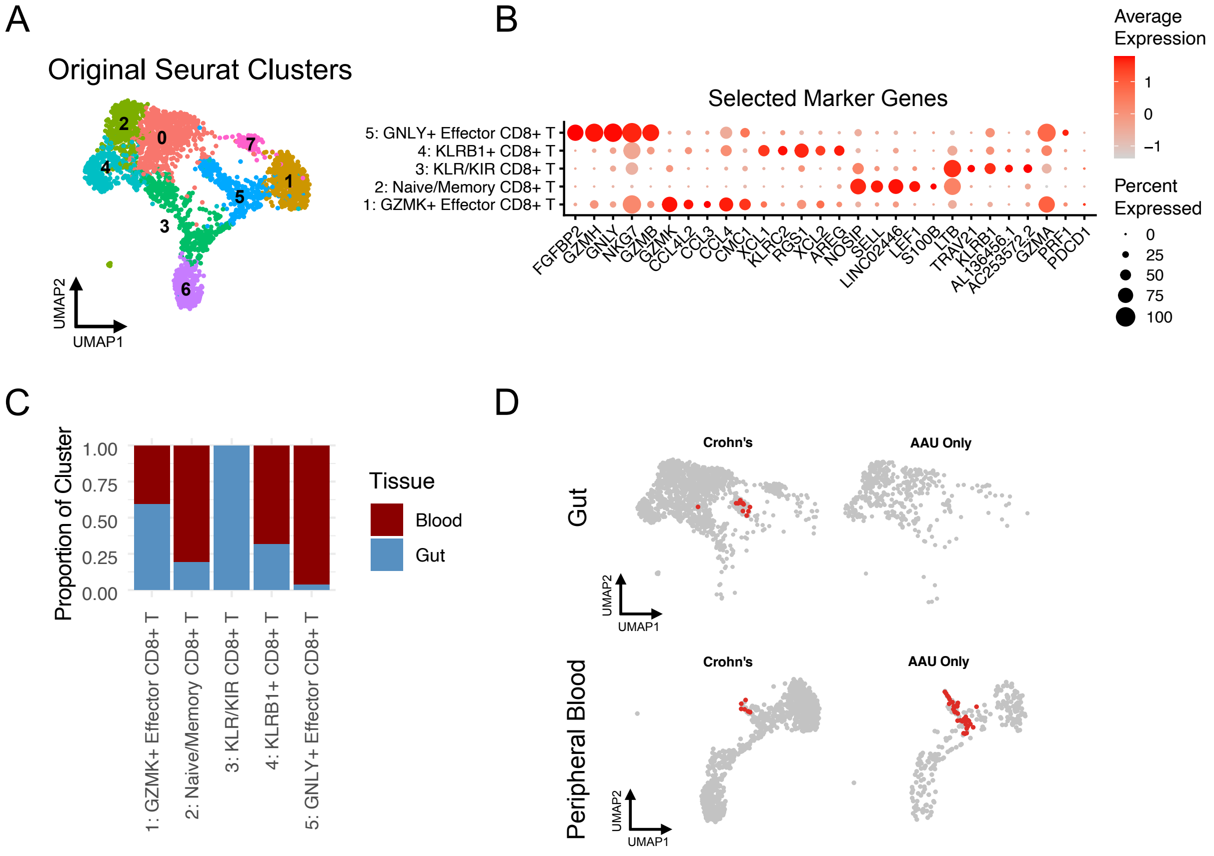


**Figure S5. (A)** UMAP plot depicting Seurat clustering of the HLA-B*27+ IBD gut/blood scRNA-seq data before cell type annotation. **(B)** Dotplot of expression values for selected marker genes, including the top 5 differentially expressed genes (DEGs) per cluster, in the HLA-B*27+ IBD gut/blood dataset. DEGs were ranked in order of descending log2FC value; all are significant (padj < 0.01). **(C)** Barplot of the proportions of blood (maroon) or gut (blue) CD8+ T cells in each cluster shown in Fig. 5B. **(D)** UMAP plots highlighting cells with pathogenic TCR motifs (red) in the HLA-B*27+ IBD gut/blood scRNA-seq data. Plots are split by tissue (rows) and IBD diagnosis (columns) for more granular visualization.

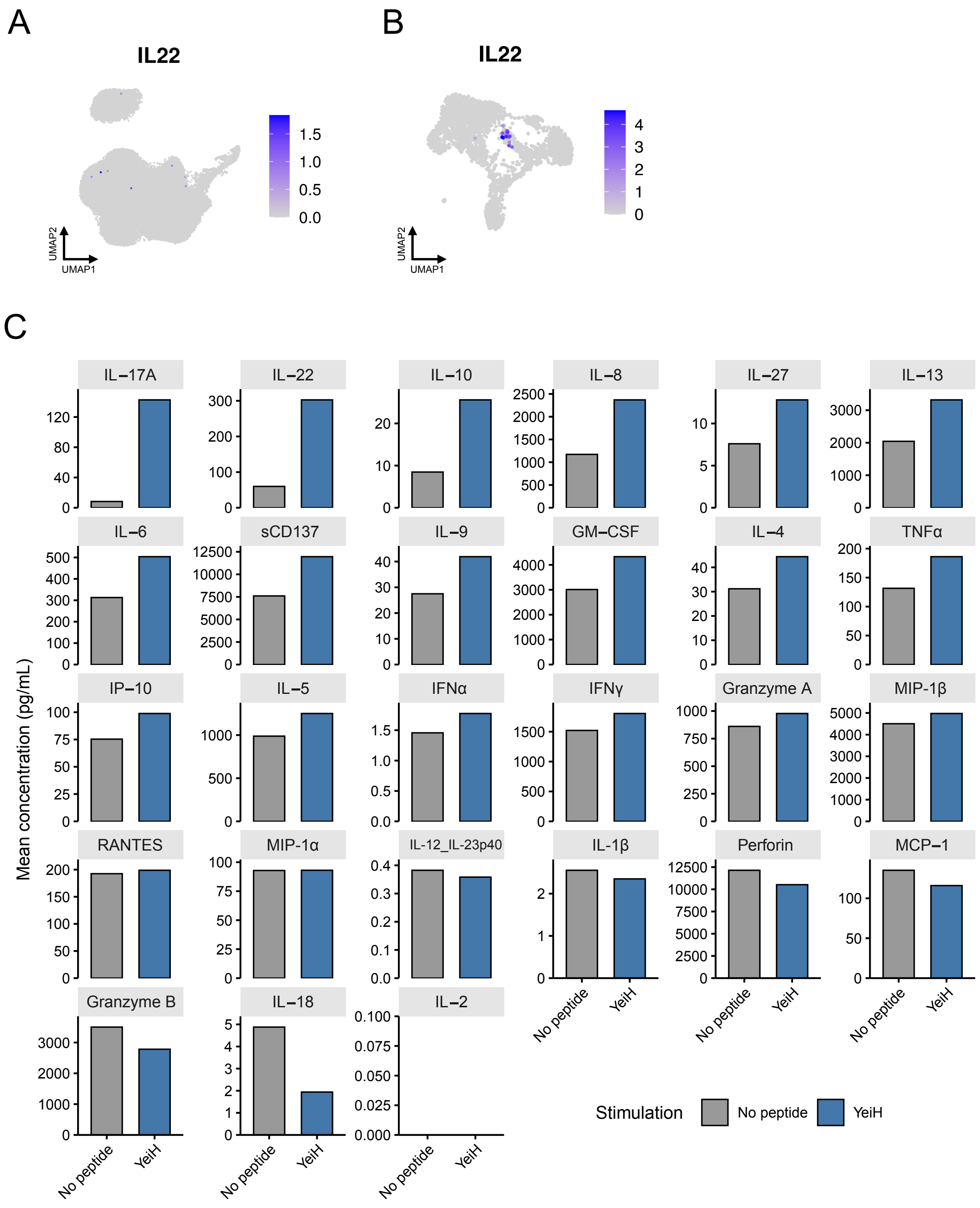
A
IL22
1.5
1.0
0.5
0.0
UMAP2
UMAP1
B
IL22
4
3
2
1
0
UMAP2
UMAP1
C
Mean concentration (pg/mL)
IL−17A
IL−22
IL−10
IL−8
IL−27
IL−13
IL−6
sCD137
IL−9
GM−CSF
IL−4
TNFα
IP−10
IL−5
IFNα
IFNγ
Granzyme A
MIP-1β
RANTES
MIP-1α
IL-12_IL-23p40
IL-1β
Perforin
MCP−1
Granzyme B
IL−18
IL−2
No peptide
YeiH
Stimulation
No peptide
YeiH

**Figure S6. (A** and **B)** Expression of *IL-22* in the public joint/eye/blood dataset (Figure 4) (A) and the HLA-B*27+ IBD gut/blood dataset (Figure 5) (B). **(C)** Cytokine production of YeiH+CD8+ T cells after stimulation with HLA-B*27+ K562 cells alone (no peptide) or pulsed with YeiH.232-240. Concentrations plotted are the mean of two biological replicates.

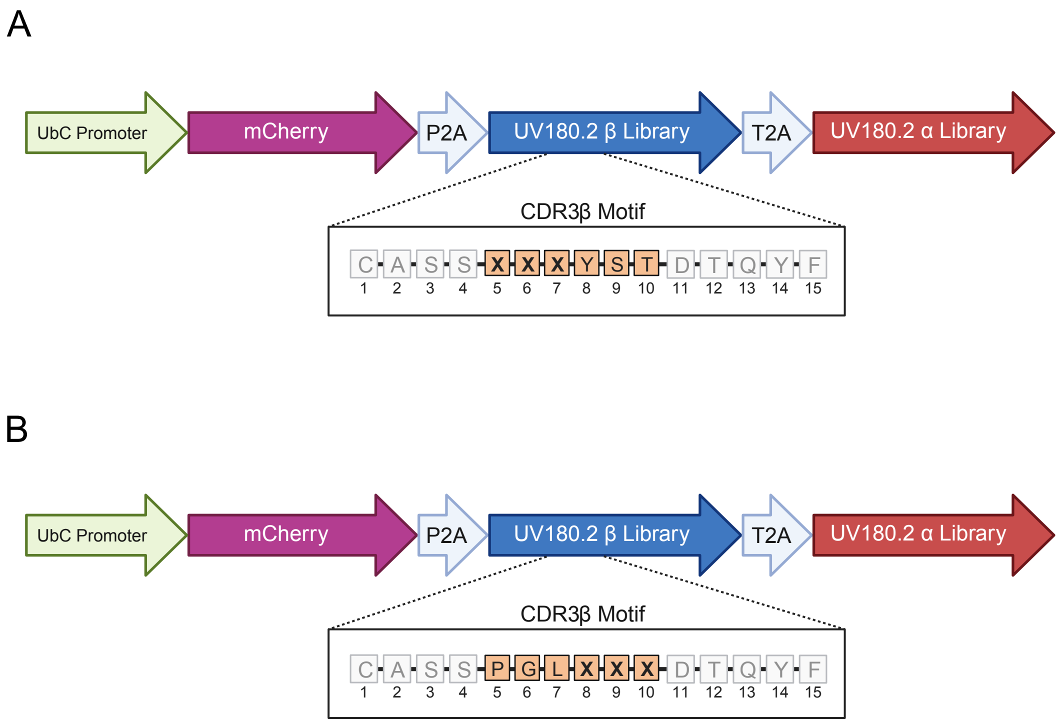


**Figure S7. (A** and **B)** Illustration of the generation of the CDR3β mutational libraries for **Figure 1**. An NNS randomization strategy was used for codon generation at CDR3β positions 5-7 (A) and CDR3β positions 8-10 (B).

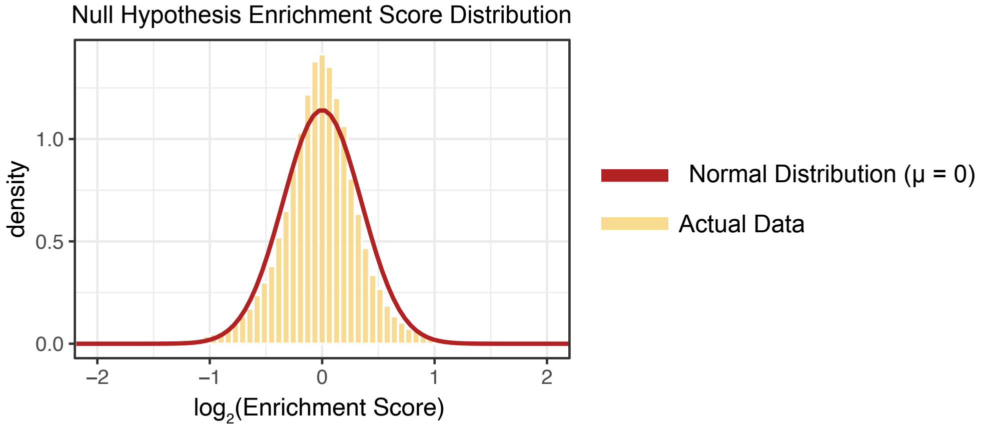


**Figure S8.** Representative plot of the null hypothesis distribution for the T-tests used in the CDR3β mutational analysis (see **Supplementary Materials and Methods**). Under the null hypothesis, each motif should be represented approximately equally in the pre- and post-selection libraries; that is, the distribution of the post-selection libraries should appear similar to the pre-selection library. To model this null hypothesis, we generated two pilot replicates of the pre-selection library, designated Replicate 2 as the "sample" and Replicate 1 as the pre-selection "control", and generated enrichment scores for Replicate 2 as described in the **Supplementary Materials and Methods.** Enrichment score values were $\log_2$-transformed and plotted. A normal distribution ($\mu = 0$, $\sigma = 0.35$) is shown in red; actual enrichment score distribution for the full library (32,723 motifs) is shown in yellow.
