## Supplemental Methods for "Refinement of the Pathogenic T Cell Receptor Motif Suggests Type 17 CD8+ T Cells Initiate HLA-B*27-associated Autoimmunity"

### SUPPLEMENTARY MATERIALS AND METHODS

*Sex as a biological variable:* Both male and female participants were enrolled in this research study and were represented in the re-analysis of public sequencing datasets.

*BLAST analysis:*

Amino acid sequences for all human TRBV gene segments were downloaded from the Swiss-Prot database (accessed through the UniProtKB website)*(24)*. Sequences encoding "probable non-functional" TRBV segments were excluded from analysis. Sequences were submitted to the NCBI Standard Protein Blast web server*(22, 23)* using the multiple sequence alignment functionality with `expect threshold` set to 1. The *TRBV9* amino acid sequence was used as the reference sequence for the BLAST search. Results and distance tree were downloaded from the BLAST web server and re-plotted in Python (v3.13.5) using the Biopython package. Phylogenetic tree plotting in Python was assisted by Claude (Anthropic; Claude Sonnet 5). All code and figure outputs were reviewed by the authors for accuracy.

*TRBV substitution:*

To determine which TRBV gene segments were supportive of pathogenicity, we modified the UV180.2 TCR (an intraocular, clonally expanded TCR from AAU, previously shown to cross-react to the YeiH.232-240 peptide as well as autoantigens)*(7, 9)*. TRBV gene segments *TRBV9*01, TRBV5-5*, *TRBV20-1, TRBV2, TRBV5-1, TRBV5-4, TRBV6-5, TRBV7-2, TRBV7-6,* and *TRBV30* were selected to replace *TRBV9*02,* the native TRBV gene segment of UV180.2. The modified TCRs were cloned into a lentiviral-based vector (pUltraHot). pUltra-hot was a gift from Malcolm

Moore (Addgene plasmid # 24130 ; http://n2t.net/addgene:24130 ; RRID:Addgene_24130). The full-length sequences of the α- and β-chains were cloned separately and transduced with pMD2G and psPAX2 packaging vectors. Lentivirus was generated for each TCR in embryonic kidney 293T cells using the Invitrogen Lipofectamine 3000 transfection kit. Cells were cultured in Opti-MEM media for 48 h, after which medium was collected. For each TCR, 1mL of lentivirus-containing media was used to infect ~500,000 CD8+ TCR-deficient Triple Parameter Reporter Jurkat cells*(21)*. After 3-4 days of expansion in RPMI complete medium containing 10% FBS, successful transduction was validated by staining for FITC anti-CD3 (eBioscience), PerCP-Cy5.5 anti-CD8a (eBioscience), TCRαb (BD Horizon), and YeiH.232-240 and analyzed on the Cytek Aurora flow cytometer (Cytek Biosciences).

HLA-B*27+ K562 cells have been previously described*(7)*. These HLA-B*27+ K562 cells were cultured with YeiH.232-240 (LRVMMLAPF), GPER1 (GQMWLLAPR), PRPF3 (TRLALIAPK), or RNASEH2B (GQVMVVAPR) as previously described*(7)*. In brief, peptides were reconstituted in 100% Dimethyl sulfoxide (DMSO). Peptides were added to 100 µl of HLA-B*27+ K562 cells to reach a 10 µM final concentration. After a 90-minute incubation period, K562 cells were washed with RPMI complete medium twice and then co-incubated with the TCR-transfected Jurkat cells (each TCR separately) for 14–18 h at 1:1 ratio. After incubation, the frequency of NFAT-GFP reporter fluorescence was measured by flow cytometry using the Cytek Aurora flow cytometer (Cytek Biosciences). Values were plotted in GraphPad PRISM.

*CDR3β mutational library generation:*

The UV180.2 CDR3β was subjected to random mutation at CDR3β amino acid positions 5-7 and 8-10, respectively. To limit the size of the libraries and minimize STOP codons, an NNS randomization strategy was used for codon generation. The UV180.2 TCRa and UV180.2 TCRb mutational libraries were cloned into the pUltraHot vector by the Genome Engineering & Stem Cell Center (GESC@MGI) at Washington University in Saint Louis, MO (**Fig. S7**). Lentivirus was generated by the HOPE Center Viral Vectors Core. Lentiviral libraries were used to transduce Jurkat cells as described above.

Jurkat libraries were expanded and sorted on CD3+mCherry+ transduced cells. Sorted Jurkats were expanded again and then co-cultured with peptide-pulsed HLA-B*27+ K562 cells overnight. Jurkat cells were then sorted on GFP+ cells as a readout for NFAT signaling and frozen for TCRbeta sequencing. If the number of sorted cells was less than 60,000 cells, untransduced Jurkats were added to the collection tube to increase the cell number to 60,000 to aid in cell pelleting during centrifugation.

Amplicon libraries were generated using a two-step PCR workflow. In the first PCR, target regions were amplified using vector-specific primers containing partial Illumina adapter tails. The forward primer tail was 5′-CACTCTTTCCCTACACGACGCTCTTCCGATCT-3′ and the reverse primer tail was 5′-GTGACTGGAGTTCAGACGTGTGCTCTTCCGATCT-3′. Primer pairs used were MS2909.YST.DS.F/MS2909.YST.DS.R and MS2908.PreYST.DS.F/MS2908.PreYST.DS.R:

MS2909.YST.DS.F: TTCCTTGGAGCTTGGGGAC

MS2909.YST.DS.R: ACAGTCAGTCTAGTTCCCGG

MS2908.PreYST.DS.F: CATATTGGAGAGGTTCTCGGC

MS2908.PreYST.DS.R: ACAGTCAGTCTAGTTCCCGG

PCR1 was performed using JumpStart REDTaq ReadyMix (Millipore Sigma, cat# P0982-800RXN). Reactions were performed with an initial 2-minute denaturation at 94°C, followed by 30 cycles of 94°C for 20 seconds, 58°C for 20 seconds, and 72°C for 40 seconds, with a final extension at 72°C for 2 minutes.

Unique dual indexes were introduced in a second PCR using 1–2 µL of PCR1 product as template. PCR2 was performed with JumpStart REDTaq Hot Start ReadyMix under the same conditions as PCR1 but for 5 cycles. Indexed PCR products were purified using DINOMAG SPRI beads (Labscoop, cat# DN9004-75ML) according to the manufacturer's recommendations. Final libraries were submitted to the Genome Technology Access Center at the McDonnell Genome Institute (GTAC@MGI) for paired-end 2 × 150 bp sequencing on an Illumina NovaSeq platform.

*CDR3β mutational analysis:*

Fastq files were obtained from GTAC@MGI and fastp (v0.20.1) was used to remove low-quality sequences. All settings were default, except for the flags `--trim_front1 15` and `--trim_front2 15`.

After fastp filtering, a custom Python read counting pipeline was used to process the filtered fastq files into read count matrices. In brief, the 5' (TCTAGT) and 3' (TATTCG) flanking sequences for each mutated region are known, and the mutated region is always 9 nucleotides long; therefore, the custom script searches for, in order, the 5’ flanking sequence, a 9-nucleotide variable sequence, and the 3’ flanking sequence. If such a sequence is found in the forward read, the reverse read is searched for the reverse complement. If the exact reverse complement is found in the reverse read, and the 9-nucleotide variable sequence is known to exist in our library, the read is counted; otherwise, the read is discarded. No mismatches are allowed. Using this protocol, approximately 90% of reads per sample were counted and mapped to known sequences in the library. Full quality control (QC) reports for the count matrix generation can be found in **Data File S7.** Code for the read count matrix generation was hand-written, with code cleanup and beautification assisted by Claude AI (Anthropic; Claude Sonnet 5). Claude was also used to generate code for the QC reports. All code and outputs were reviewed by the authors for accuracy.

Post-selection motif enrichment analysis followed a custom R pipeline based on deep mutational sequencing methods*(7, 43)*, as follows.

1. Within each sample, all count values were divided by total counts per sample, then multiplied by a factor of 1,000,000 to achieve a “counts per million” (CPM) normalized matrix.
2. A pseudocount of 1 was added to the entire matrix.
3. For each post-selection sample, an “enrichment score” was calculated for each nucleotide-level variable sequence. Enrichment score was calculated as the ratio of the sequence’s value in the post-selection sample to the sequence’s value in the pre-selection starting library sample.
4. The enrichment score matrix was log2-transformed.

*Note about further statistical testing:* We produced two technical replicates for each condition (pre-selection, GPER1 selection, PRPF3 selection, RNASEH2B selection, and YeiH selection). Given that our mutational library was randomly generated at the nucleotide level, most (76%) amino acid motifs in our library were represented by at least two degenerate codons. We reasoned that true hits should show consistent directional selection across all degenerate codons encoding the same motif. We therefore treated each degenerate codon as an independent replicate in the step below.

5. Under our null hypothesis, motif abundance should be unchanged before and after selection (that is, the enrichment score for each motif should equal 1, and the log-transformed enrichment score should equal zero.) Therefore, for each amino acid sequence present in our log-transformed enrichment matrix, statistical significance was measured using a one-sample t-test with mu = 0.

   *Note:* A T-test was chosen because the ratio of any two pre-selection library replicates followed an approximately log-normal distribution (see **Fig. S8**).

6. P-values were adjusted using Benjamini-Hochberg (FDR) correction. Motifs were considered statistically significant if they met the following criteria: adjusted p-value < 0.05, |log2FC| > 1.

Volcano plots for the enrichment statistics were plotted in R using the `ggplot2` package (v3.5.2). Positively-selected motifs passing significance cutoffs were plotted using the `VennDiagram` package (v1.8.2).

Cross-reactive motifs were defined as any motifs which were significantly positively selected by YeiH as well as 1 or more human autoantigens (GPER1, PRPF3, and/or RNASEH2B). Cross-reactive motifs were plotted as a SeqLogo plot using the `ggseqlogo` package (v0.2.2), a heatmap of amino acid proportions per position using the `pheatmap` package (v1.0.12), and an alluvial plot using the `ggalluvial` package (v0.12.5). Plotting code was assisted by Claude AI (Anthropic; Claude Sonnet 5). All code and outputs were reviewed by the authors for accuracy. All analysis code is available on GitHub (see **Data, Code, and Materials Availability**).

*TIRTL-seq:*

*Biospecimen collection:*

Peripheral blood samples were obtained by venipuncture and collected into EDTA tubes. Peripheral blood mononuclear cells (PBMCs) were isolated by Ficoll-Hypaque density gradient centrifugation. PBMCs were cryopreserved in fetal bovine serum (FBS) containing 10% dimethyl sulfoxide (DMSO) and stored at -140ºC until analysis.*(9)*

*Sample Collection and Library Preparation:*

We obtained peripheral blood from 8 HLA-B*27+ healthy controls and 6 HLA-B*27+ patients (2 with axSpA only, 4 with axSpA and AAU). TIRTL-seq was performed as previously described*(25)*. Briefly, cryopreserved PBMCs were thawed, washed in RPMI 1640 media supplemented with L-glutamine, penicillin/streptomycin, and 10% fetal bovine serum, and resuspended in calcium- and magnesium-free DPBS (Thermo Scientific). $CD8^+$ and $CD4^+$ T cells were isolated by positive selection using EasySep Human CD8 Positive Selection Kit II (STEMCELL Technologies, Cat# 17853) and EasySep Release Human CD4 Positive Selection

Kit (STEMCELL Technologies, Cat# 17752), respectively, according to the manufacturer's instructions.

Cell lysis and cDNA synthesis were performed in 384-well plates preloaded with Vapor-Lock (Qiagen). Cell suspensions were combined with an RT/lysis master mix containing Maxima H Minus Reverse Transcriptase (4 U/µL), Triton X-100 (0.1%), RNasin (1 U/µL), dNTPs (0.5 mM each), DTT (5 mM), random hexamers (2.5 ng/µL), and oligo(dT) (2.5 µM) using a MANTIS liquid dispenser (Formulatrix). Plates were incubated at 42°C for 5 min, 25°C for 10 min, 50°C for 60 min, and 94°C for 5 min.

TCRα and TCRβ cDNA were multiplex-amplified (PCR I) using KAPA2G Fast Multiplex Mix (Roche), a pool of 88 V-segment forward primers (2.25 µM each), and plate-barcoded TRAC/TRBC reverse primers (10 µM each). All primers (found in the Supplementary Data 1 file of the original TIRTL-seq manuscript) were synthesized by Integrated DNA Technologies*(25)*. PCR I was performed at 95°C for 3 min, followed by 20 cycles of 95°C for 15 s, 59°C for 30 s, and 72°C for 1 min, with a final extension at 72°C for 5 min.

To generate the libraries, PCR I products were diluted 1:20 and indexed (PCR II) using Q5 Hot Start High-Fidelity DNA Polymerase (NEB), Q5 Reaction Buffer, dNTPs (0.2 mM each), and unique dual-indexed i5/i7 primers (1 µM). PCR II consisted of 15 cycles of 98°C for 10 s, 58°C for 10 s, and 72°C for 50 s, followed by a 2-min extension at 72°C. Finally, indexed libraries were pooled, purified using AMPure XP beads (Beckman Coulter), and quantified by TapeStation using High Sensitivity D5000 ScreenTape (Agilent) and High Sensitivity D5000 Reagents (Agilent).

Libraries were sequenced on an Illumina NovaSeq platform targeting 150,000,000 reads per library.

*TCR repertoire processing*:

Raw FASTQ files were processed using an implementation of the TIRTL-seq workflow (cite). Wells sequenced across multiple flow cells were concatenated and verified before alignment. Reads were aligned and clonotypes assembled with MiXCR (v4.7.0)*(44)* (reference library repseqio. v5.1) using the analyze-generic-amplicon preset with the human reference in RNA mode, a floating left alignment boundary, a floating right boundary anchored on the C segment, and clonotype assembly by CDR3*(44)*. Well barcodes were parsed from an 8- nucleotide tag on read 1 and demultiplexed to plates using a plate index sheet. Wells failing any of four quality criteria were flagged: fewer than 5,000 aligned reads, a TRA or TRB read fraction above 0.85, or an alignment rate below 0.50. Per-well clonotype tables were mapped to 384-well plate coordinates, restricted to plate identifiers used in the experiment to exclude index-hopping artifacts, and concatenated into per-plate TRA and TRB dictionaries carrying a well identifier.

*Pairing and quantification*:

Pairing was performed per plate on wells whose clonotype counts exceeded 50% of the plate mean in both chains, using sparse clone-by-well read-fraction matrices restricted to clonotypes observed in more than two wells. Alpha-beta pairs were identified with MAD-HYPE*(45)* and the T-Shell correlation method as described in Pogorelyy et al.*(25)*, both executed on a GPU through an MLX v0.31.2 backend under Python 3.13.5. Pairs from both methods were retained when they overlapped in more than two wells and had a combined chain-loss fraction below 0.05. T-

Shell calls additionally required an adjusted p-value below 1e-10. Surviving pairs were annotated with CDR3 amino acid sequences and modal V, D, J, and C gene assignments. Plate- level clone frequencies were joined from the bulk dictionaries, with QC- flagged wells excluded from the denominator. The final paired matrix represented the union of all alpha-beta sequences that were found via either algorithm (T-SHELL or MAD-HYPE). Single-chain pseudobulk tables were generated in parallel by aggregating each dictionary over unique CDR3 nucleotide sequences. Well occupancy, aggregate read-count statistics, per-well read-count statistics, modal V and J genes, and read fraction were reported in the column format expected by TIRTLtools.

Percentages of pathogenic T-cell receptors (TCRs) were quantified per subject under three progressively more specific criteria, as follows. *Criteria 1*: CDR3β sequence matching the broadened CDR3β motif (A/G/D/S/E/Q/R in position 6, F/R/L/Y/H/V/I/T/S/M in position 7, Y/F in position 8, S in position 9, and T in position 10) ("Criteria 1"). *Criteria 2*: Criteria 1 plus TRBV gene segment matching *TRBV9, TRBV5-5,* or *TRBV5-4*. *Criteria 3*: Criteria 2 plus TRAV gene segment matching *TRAV21*. We applied Criteria 1 and 2 to all available TCRβ sequence data, then applied Criteria 3 to a filtered version of the dataset where the only allowed sequences were alpha-beta pairings with exactly one alpha and one beta sequence per cell. Data were plotted as dotplots in R; mean and standard deviation were calculated and overlaid onto the dotplots. Two-sided Wilcoxon rank-sum (Mann-Whitney U) tests were performed to determine the statistical significance of the difference between the healthy and disease distributions.

Sensitivity and specificity of each test was assessed using the `pROC` package*(46)* (v1.18.5). We first called the `pROC::roc` function, setting disease status as the outcome, pathogenic TCR percentage as the predictor, and axSpA/AAU samples as the positive class. Then, to obtain sensitivity and specificity metrics using our "presence/absence" cutoff, we called the `pROC::coords` function and set the threshold to 1e-100 (chosen as an arbitrarily low nonzero value).

*YeiH+CD8+ T cell sort scRNA-seq, scTCR-seq, and data analysis*

*Single Cell Sample Preparation:* CD8+ T cells were isolated from peripheral blood mononuclear cells (PBMCs) from five healthy controls and four participants with AAU and/or axSpA via EasySep™ Human CD8+ T Cell Isolation Kit (StemCell Technologies, Vancouver, Canada). Cells were labeled with TotalSeq-C hashtags (Biolegend, San Diego, California), stained with antibodies for CD3 FITC (Invitrogen #11-0037-42), CD8 PerCP-Cy5.5 (Invitrogen #45-0088-42), and HLA-B*27(YeiH.232-240) tetramers labeled with either PE or APC. Cells were sorted for CD3+ CD8+ and YeiH APC/PE double-positive cells on BD FACS Aria III (BD Biosciences, Franklin Lakes, New Jersey). The 16,000 sorted cells were combined with 44,000 "bystander" PBMCs from an unrelated sample that was labeled with a separate hashtag for centrifugation and cell pellet visualization. The entire 60,000 cells were re-suspended in 37.5ul for super-loading of the 10x Chromium Controller (10x Genomics).

*Library Preparation:* Single-Cell 5′ Gene Expression cDNA libraries were generated per the 10x Genomics Chromium Single-Cell 5′ Library and Gel Bead Kit v2 and the 10x Chromium Controller platform for microdroplet-based, single-cell barcoding, by the Genome Technology Access Center at the McDonnell Genome Institute (GTAC@MGI, Washington University in St.

Louis). T cell enrichment libraries were generated per the Chromium Single-Cell V(D)J Enrichment Kits for Human T cell (10× Genomics) using the same input samples. All libraries were sequenced at GTAC@MGI on the NovaSeq Sequencing System (Illumina).

*Data Preprocessing:*

Preprocessing of the scRNA-seq and scTCR-seq FASTQ files was performed using the 10X Genomics CellRanger software (v8.0.1) and the GRCh38 human genome reference. The command 'cellranger multi' was used to align the paired scRNA-seq and scTCR-seq FASTQ files to the reference genome and quantify the number of reads per gene. Hashtag oligo (HTO)-based antibody hashtags were specified as an additional feature input during CellRanger alignment.

After CellRanger processing, the filtered scRNA-seq and HTO count matrices were imported into R (v4.4.0) and manually cleaned using the `Seurat` package*(47, 48)* (v4.4.0). Cells were removed if they displayed a high mitochondrial gene percentage (indicating low quality), unusually low gene count, or unusually high unique molecular identifier (UMI) count (indicating probable doublets). These quality cutoffs are detailed in **Data File S9**. After quality filtering, we obtained gene expression data for 2,652 cells across the nine samples (median: 239 cells per participant).

*Sample Demultiplexing:*

HTO-based sample demultiplexing was performed in R on the HTO matrix using the `demuxmix` package (v1.0.0) and the default Demuxmix pipeline*(49)* (https://www.bioconductor.org/packages/release/bioc/vignettes/demuxmix/inst/doc/demuxmix.ht ml). After demultiplexing, cells belonging to the “bystander” sample were removed from further analysis.

*Clustering Analysis:*

Filtered and preprocessed gene matrices were clustered using Seurat, following the default pipeline with minor modifications*(47, 48)*. In brief, the matrices were log-normalized using the `NormalizeData` function, and the top 2000 variable genes were identified using the `FindVariableFeatures` function. To prevent TCR-specific genes from driving clustering, the function `quietTCRgenes` from the `Trex` package (v0.99.11)*(50)* was called on the Seurat objects prior to scaling with Seurat’s `ScaleData` function. The effects of sex were regressed out of the clustering during the `ScaleData` step. Principal components were identified using `RunPCA` after scaling. Downstream K-Nearest Neighbors analysis, cluster analysis, and UMAP dimensionality reduction were run using the Seurat functions `FindNeighbors`, `FindClusters`, and `RunUMAP`, respectively. Parameters for these functions can be found in **Data File S9**. Once the data was clustered, any contaminant (non-CD8+ T cell) clusters (those that did not co-express CD45, CD3, and CD8) were removed, and the object was re-clustered from the FindVariableFeatures step to the RunUMAP step.

Clusters were manually annotated with cell type labels according to the expression of marker genes. $LEF1^{+}TCF7^{+}CCR7^{+}SELL^{+}CD8^{+}$ T cells were annotated as naive/T central memory. $GZMA^{int}NKG7^{int}LEF1^{int}CD8^{+}$ T cells were labeled as early effector CD8+ T cells. $LEF1^{-}GZMA^{+}GZMK^{+}PRF1^{+}GNLY^{+}NKG7^{+}CD8^{+}$ T cells were labeled as “late” or terminally differentiated effector cells. *LEF1*$^{-}$*GZMA*$^{+}$*NKG7*$^{+}$*KLRB1*$^{+}$*CD8*$^{+}$ T cells were labeled as KLRB1+ effector cells. Cluster percentages per sample were plotted using GraphPad PRISM.

*Differential Gene Expression Analysis:*

Differentially expressed genes (DEGs) were identified using the MAST*(51)* algorithm; this algorithm was called through the Seurat `FindMarkers` wrapper function. Genes were considered differentially expressed if they passed a |log2FC| threshold of 0.5 or higher and an adjusted p-value of 0.05 or lower. Volcano plots for DEGs were constructed using the `draw.volcanoPlot` function from the package `NetBID2` (v2.0.3)*(52)*.

*Pathway Analysis:*

Signatures for pathway analysis were derived from multiple public resources, as follows. Canonical Pathways signatures (c2.cp.v2023.2.Hs.symbols) and ImmuneSigDB signatures (c7.immunesigdb.v2023.2.Hs.symbols) were downloaded directly from the MSigDB. Th17 signatures were taken from the Supplementary Data of Ramesh et al.*(37)*. CD161 signatures were generated as follows: processed microarray data from CD161$^{neg}$, CD161$^{int}$, and CD161$^{high}$ CD8+ T cells were downloaded from the GEO database (GSE62096) and re-analyzed in R using the `limma` package (v3.60.2) as previously described*(37, 53)*. Genes passing a |logFC| cutoff > 1 and an adjusted p-value cutoff < 0.05 were considered differentially expressed. The "CD161int signature" shown in **Fig. 3J** is the positively differentially expressed genes in the CD161$^{int}$ cells compared to the CD161$^{neg}$ cells. The MAIT cell cluster DEGs were taken from the differential gene expression analysis detailed in the "Joint, Eye, and Blood Public Dataset scRNA-seq, scTCR-seq, and Data Analysis" of this Methods section.

To determine the significance of overlap between the pathogenic gene signature and public gene signatures of interest, Fisher's Exact Test was performed on the Canonical Pathways signatures, the Th17 signatures, the CD161 signatures, and the MAIT cluster DEGs using the `funcEnrich.Fisher` function from the R package `NetBID2`*(52)*. Multiple testing p-value correction was performed using the Benjamini-Hochberg method.

To score each cell in the dataset for the activity of the public signatures, we used the `AUCell` package*(30)* (v1.26.0). We extracted the count matrix from the cleaned Seurat object, then used the `AUCell_run` function to build gene expression rankings within each cell and calculate area under the curve (AUC) for the recovery curve of each gene set. AUC scores for each signature were added back to the Seurat object using Seurat's `AddMetaData` function and plotted using Seurat's `FeaturePlot` function.

*TCR Analysis:*

After alignment, the "filtered contig annotation" matrices from the CellRanger output were used for downstream scTCR-seq analysis. TCR sequences were analyzed in R using the `scRepertoire` package, following the default scRepertoire pipeline*(54)* (v2.0.0). Paired TCR sequencing data were available for 2,520 (95%) of the cells in the dataset. TCRs matching Criteria 3 (from the "TIRTLseq" section of the Methods) were considered pathogenic.

*axSpA and PsA Public Bulk TCRβ Sequencing Analysis*

Fastq files were downloaded from Komech et al.*(8)* through ArrayExpress accession E-MTAB-11498. MiXCR (v4.7.0) was used to align and assemble the reads and output tables of the assembled clonotypes*(44)*. As UMI data did not appear to be available in the fastq reads, MixCR's `analyze` command was used on the `generic-amplicon` setting with flags `--species hsa`, `--rna`, `--rigid-left-alignment-boundary`, `--floating-right-alignment-boundary C`, and `--assemble-clonotypes-by CDR3`. MiXCR outputs were read into R using the `immunarch` package (v0.10.3.9002)*(55)* and the percentage of reads containing both the broadened pathogenic CDR3β motif and an allowed TRBV gene segment (*TRBV9, TRBV5-5,* or *TRBV5-4*) was calculated. These percentages were plotted in PRISM (v11.0.2).

*Joint, Eye, and Blood Public Dataset scRNA-seq, scTCR-seq, and Data Analysis*

For the axSpA and PsA datasets, fastq files were downloaded from the SRA database: SRP403458 (Deschler et al.), SRP405490 (Povoleri et al.), SRP561214 (Tang et al.)*(19, 31, 32)*. For the AAU dataset (Paley et al.), fastq files were obtained upon request from the authors*(9)*.

*Data Preprocessing:* All fastq files were processed using the 10X Genomics CellRanger software (v8.0.1) and the GRCh38 human genome reference, following the same procedures detailed in "Data Preprocessing" above ("YeiH+CD8+ T cell sort scRNA-seq, scTCR-seq, and data analysis"). Because each dataset displayed a different average sequencing depth and mitochondrial percentage, filtering was performed on each dataset separately. Cutoffs used for each dataset can be found in **Data File S9.** Sample SRR22025519 (Deschler et al.) displayed low overall quality and was removed from further analysis.

*Sample Demultiplexing:* As the AAU dataset samples were multiplexed with antibody hashtagging, sample demultiplexing was performed as detailed in "Sample Demultiplexing", above.

*Clustering Analysis:* Filtered and preprocessed gene matrices were clustered using Seurat v5.2.0*(56)*. In brief, the matrices were log-normalized using the `NormalizeData` function, and the top 2000 variable genes were identified using the `FindVariableFeatures` function. Data was scaled using `ScaleData` with default parameters, and principal components were identified using `RunPCA`. Because there was a batch effect between samples and datasets, sample-level integration was performed using the Harmony package*(57)* (v1.2.3) through the Seurat v5 wrapper function `IntegrateLayers`. Downstream K-Nearest Neighbors analysis, clustering, and UMAP dimensionality reduction were run as described above with the parameters found in **Data File S9**. Contaminant clusters (those that did not co-express CD45, CD3, and CD8) were removed.

Clusters were manually annotated according to their expression of marker genes. Naive CD8+ T cells were defined as those that expressed *LEF1, CCR7,* and *TCF7* in the absence of effector genes (*GNLY, PRF1, GZMA, GZMB, NKG7*). Cycling CD8+ T cells were defined as those that expressed cell cycle-related genes (*MKI67, STMN1, PCNA*). The MAIT cell cluster was identified by high expression of *TRAV1-2* and *KLRB1.* All other conventional CD8+ T cell clusters were labeled in accordance with their marker gene expression (*GZMK* for the GZMK+CD8+ T cluster; *KLRB1* for the KLRB1+CD8+ T cluster; *GNLY* for the GNLY+CD8+ T cluster; and *KIR3DL1*, *KIR2DL4, KLRC2*, and *KLRC3* for the KIR/KLR+CD8+ T cluster).

*Differential Gene Expression Analysis:*

Given the batch effect and large cell numbers present in the dataset, DEG analysis was performed using a pseudobulk approach to minimize false positives. The `DESeq2` package (v1.44.0) was used as described*(58)* (https://bioconductor.org/packages/devel/bioc/vignettes/DESeq2/inst/doc/DESeq2.html). Briefly, the count matrix was pseudobulked by subject, tissue, and cluster, then filtered to remove genes that were expressed (defined as 10 counts or more) in less than 3 pseudo-samples. The DESeq2 object was built with the formula `~0 + subject + tissue + cluster`, and the DESeq2 algorithm was run using the "local" fit type. For each cell type, a contrast vector was set such that the mean expression of the genes in that cell type were compared to the mean expression of those same genes in all the other cell types in the dataset. Genes that passed an adjusted p-value cutoff of 0.01 and a log2FC cutoff of 1 were considered differentially expressed.

*Pathway Analysis:*

Activity of the pathogenic gene signature (**Fig. 3I**) was calculated in the dataset using AUCell, as described in "Pathway Analysis," above ("YeiH+CD8+ T cell sort scRNA-seq, scTCR-seq, and data analysis"). Fisher's Exact test comparing the MAIT cell DEGs to the pathogenic signature was performed as part of the Fisher's Exact Tests described in "Pathway Analysis," above ("YeiH+CD8+ T cell sort scRNA-seq, scTCR-seq, and data analysis").

*TCR Analysis:*

TCR analysis was carried out using the `scRepertoire` package, as described in "TCR Analysis," above ("YeiH+CD8+ T cell sort scRNA-seq, scTCR-seq, and data analysis").

For the barplots of TCR features based on gene signature expression scores, we first subsetted the dataset to include only the conventional CD8+ T cell clusters (all clusters except for the MAIT cluster). We then calculated the mean AUCell pathogenic gene signature score for each clonotype in the dataset ("Pathway Analysis," above). Clonotypes were ranked highest to lowest based on their average score and assigned a percentile. Clonotypes with the specified TCR features of interest (pathogenic CDR3β sequence or presence of the *TRBV9/5-5/5-4* gene segment) were highlighted in the barplots. For the TRAV21+ conventional CD8+ T cell barplots, the same procedure was repeated after narrowing the dataset to only those conventional CD8+ cells whose TCRs contained the *TRAV21* gene segment. For the control geneset, 20 genes (the same number of genes in the pathogenic gene signature) were randomly selected from the genes present in the dataset. The control geneset was scored in each cell using AUCell (see "Pathway Analysis") and the same barplot procedures detailed above were repeated using the control geneset in place of the pathogenic geneset. All barplots were generated using the `ggplot2` package for R.

*Identification of MAIT cell TCRs:* PBMCs from five healthy individuals were labelled with TotalSeq C Hashtag antibodies, combined, and stained with CD3, CD161, and MR1(5-OP-RU) PE tetramer (NIH Tetramer core, Atlanta, GA). Cells were sorted for $CD3^{+}CD161^{hi}MR1^{+}$ T cells and subjected to single-cell TCR sequencing and analysis as described above. The TCR sequencing information from the eye/joint/blood public datasets was then searched for sequences exactly matching the ones from the MAIT cell sort. Cells with TCR sequences matching the MAIT cell sort were annotated as MAIT cells.

*Joint, Eye, and Blood TCR Reactivity Validation*

Twenty clonotypes were selected from Cluster 6 of the joint, eye, and blood scRNA-seq + scTCR-seq dataset (**Fig. 4D**). To ensure we were sampling evenly from the cluster, we stratified the clonotypes in the cluster into four quartiles based on their average pathogenic gene signature score and chose five clonotypes from each quartile. The chosen clonotype TCRs were cloned into a lentiviral-based vector (pUltraHot) and expressed in CD8+ TCR-deficient Triple Parameter Reporter Jurkat cells as detailed above ("TRBV Substitution"). NFAT-GFP fluorescence was quantified on the Cytek Aurora flow cytometer (Cytek Biosciences). Values were plotted in GraphPad PRISM.

*HLA-B*27+ Gut and Blood Data Collection*

*Recruitment of HLA-B*27+ Participants with Inflammatory Bowel Disease:*

A total of 44,078 electronic medical records were searched to identify 51 HLA-B*27+ individuals with a diagnosis of inflammatory bowel disease (IBD), including ulcerative colitis, Crohn's disease, and microscopic colitis.

In parallel, 992 DNA samples from participants with IBD were obtained through the Digestive Disease Research Core Center (DDRCC) at Washington University in St. Louis and screened for HLA-B*27+ by polymerase chain reaction (PCR) as previously described. An additional 62 HLA-B*27+ individuals were identified through PCR-based screening.

The two populations were merged to generate a cohort of 113 HLA-B*27+ individuals. Following secondary chart review, 14 individuals were excluded: 12 deceased patients, one duplicate identified across datasets, and one patient who met pre-specified exclusion criteria (active cancer on chemotherapy). Of the remaining eligible individuals contacted by mailed invitation, 20 responded and intestinal biopsies could be collected from four individuals undergoing a clinically indicated colonoscopy. Two participants provided paired colon and ileum biopsies; the remaining two only provided colonic biopsies. Among the four participants, two had Crohn's disease alone and were HLA-B*27+. One had Crohn's disease with HLA-B*27+ acute anterior uveitis (AAU). One biopsy sample was subsequently excluded during quality-control HLA-B27 re-typing, leaving three samples from the original cohort for analysis. An additional participant with HLA-B*27+ AAU and a family history of ulcerative colitis was recruited prior to a diagnostic colonoscopy, which ruled out IBD.

*Collection of Biospecimens for IBD Participants:*

Peripheral blood was collected from HLA-B*27+ IBD participants by venipuncture immediately before colonoscopy and PBMCs were isolated as described above. During colonoscopy, sharp-toothed biopsy forceps were used to obtain tissue samples from the colon and, when accessible, the terminal ileum. After each pass, forceps were rinsed in a labeled conical tube containing complete RPMI (RPMI-1640 supplemented with 10% FBS) to release the tissue. The target collection was 12 colonic biopsies (two each from the ascending colon, transverse colon, descending colon, cecum, and rectum) and 4 ileal biopsies. Collected samples were transported on wet ice (4°C) to the research laboratory for same-day cryopreservation.

Using sterile forceps, tissue pieces were transferred into labeled, de-identified cryovials containing 500 µL of 100% FBS. Once each biopsy type was transferred into 500 µL FBS, an equal volume (500 µL) of 2× freezing medium (20% DMSO in FBS, yielding a final concentration of 10% DMSO in FBS) was added dropwise to each cryovial with gentle mixing. Cryovials were immediately placed in an isopropanol-filled controlled-rate freezing container (Mr. Frosty; Thermo Fisher Scientific) and stored at -80°C overnight, then transferred to liquid nitrogen for long-term storage.

*Intestinal Biopsy Enzymatic Digestion:*

A pre-enzymatic digestion medium was prepared by supplementing RPMI-1640 with 2% heat-inactivated FBS, 10 mM HEPES, 100 U/mL penicillin, 100 µg/mL streptomycin, and 50 µg/mL gentamicin. Immediately prior to use, a complete enzymatic digestion mix was assembled by adding Liberase TH (100 µg/mL final concentration; Roche) and DNase I (100 µg/mL final concentration; Sigma-Aldrich) to the pre-enzymatic medium. Both enzyme stocks were reconstituted to 1 mg/mL in sterile water and stored as single-use aliquots at -20°C. The complete digestion mix (5 mL per sample) was pre-warmed to 37°C in a bead bath immediately before use.

Cryopreserved intestinal biopsy vials were thawed and immediately transferred into 50 mL conical tubes containing 30 mL of ice-cold PBS. Samples were washed twice in 30 mL ice-cold PBS with gentle inversion and careful vacuum aspiration between washes to minimize tissue disruption. Following the final wash, 5 mL of pre-warmed enzymatic digestion mix was added per sample, and tubes were incubated on a shaker at 250 rpm for 30 minutes at 37°C. Enzymatic dissociation was terminated by addition of 1 mL 100% FBS and 80 µL of 0.5 M EDTA (pH 8.0) per sample, followed by a 5-minute rest on ice.

*Processing of Digested Biopsy Samples:*

Following quenching, samples were gently triturated using a 5 mL serological pipette to achieve mechanical dissociation of residual tissue fragments. Cell suspensions were passed through 70 µm cell strainers into fresh 50 mL conical tubes. Strained suspensions were washed twice with ice-cold PBS and centrifuged at 400 × g for 5 minutes at 4°C.

For viability discrimination, cells were resuspended at $1 \times 10^6$ cells per 100 µL in PBS containing a 1:1,000 dilution of Fixable Viability Dye eFluor 780 (eBioscience) and incubated for 20 minutes on ice, protected from light. Cells were then washed once with 1 mL sort buffer (2% FBS in PBS; centrifugation at 400 × g for 5 min) and resuspended at $1 \times 10^6$ cells per 100 µL in sort buffer prior to hashtag labeling.

Blood CD8+T cells were isolated from paired PBMC samples via EasySep selection as described above. The enriched CD8+ T cells and intestinal cells were labeled with TotalSeq-C hashtags as described above.

Intestinal and blood CD8+ T cells were stained with Fixable Viability Dye eF780 (eBioscience), surface antibodies (antibody reagents provided in **Data File S8**), and barcoded PE YeiH tetramer (to identify YeiH-binding gut cells among the bulk CD8+ T cells in downstream analysis). Blood CD8+ T cells were additionally stained with APC YeiH tetramer.

*Sorting for Gut CD8+ T Cells and Blood YeiH-binding CD8+ T Cells:*

The sorted gut cells were gated on EpCAM-, CD45+, CD19-, TCR Va7.2-, TCR Va24-J18-, TCR gamma/delta-, CD3+, and CD8+. The sorted blood CD8+ T cells were gated on EpCAM-, CD45+, CD19-, TCR Va7.2-, TCR Va24-J18-, TCR gamma/delta-, CD3+, CD8+, and YeiH+. Cells were sorted into FBS.

Sorting was divided across two batches. On the first sort, 10,605 gut CD8+ T cells were collected. Due to operator error of the sorter, 2482 bulk blood CD8+ T cells were collected; after correction, 447 blood YeiH+ CD8+ T cells were collected. On the second sort, 11,170 gut CD8+ T cells and 1725 blood YeiH+CD8+ T cells were collected. Cells were subjected to single-cell RNA and TCR sequencing as described above.

*HLA-B*27+ Gut and Blood scRNA-seq + scTCR-seq Analysis*

*Data preprocessing and sample demultiplexing:* Performed as described above ("YeiH+CD8+ T cell sort scRNA-seq, scTCR-seq, and data analysis").

*Clustering Analysis:* Filtered and preprocessed gene matrices were clustered using Seurat*(47, 48)* (v4.4.0). In brief, the matrices were log-normalized using the `NormalizeData` function, and the top 2000 variable genes were identified using the `FindVariableFeatures` function. Data was scaled using `ScaleData` with default parameters, and principal components were identified using `RunPCA`. After principal component identification, samples were integrated by patient using the `RunHarmony` command from the `harmony` package*(57)* (v1.2.1). Downstream K-Nearest Neighbors analysis, cluster analysis, and UMAP dimensionality reduction were run on the Harmony embeddings using the Seurat functions `FindNeighbors`, `FindClusters`, and `RunUMAP`, respectively. Parameters for these functions can be found in **Data File S9**. Once the data was clustered, any contaminant (non-CD8+ T cell) clusters (those that did not co-express CD45, CD3, and CD8) were removed, and the object was re-clustered from the FindVariableFeatures step to the RunUMAP step. Clusters were annotated based on canonical marker gene expression, as described in the "Clustering Analysis" sections, above.

*Differential Gene Expression Analysis:* DEG analysis was performed using the MAST algorithm*(51)* through the Seurat `FindMarkers` wrapper function, as described above ("YeiH+CD8+ T cell sort scRNA-seq, scTCR-seq, and data analysis").

*Pathway Analysis:* AUCell*(30)* analysis was performed to score the pathogenic gene signature in each individual cell in the dataset, as described above ("Joint, Eye, and Blood Public Dataset scRNA-seq, scTCR-seq, and Data Analysis"). GraphPad PRISM was used to plot the mean pathogenic signature scores in Cluster 4 of the dataset.

*TCR Analysis:* TCR data processing and analysis were performed as described ("YeiH+CD8+ T cell sort scRNA-seq, scTCR-seq, and data analysis"). After quality filtering, 2,517 (80%) of the cells in the dataset had paired scTCR-seq information.

*YeiH+CD8+ T Cell Cytokine Panel*

*Enrichment of CD8+ T cells from Peripheral Blood Mononuclear Cells (PBMCs):* CD8+ T cells were enriched from bulk PBMCs using negative selection via magnetic enrichment as described above.

*YeiH Peptide Stimulation of CD8+ T Cells:*

HLA-B*27+ K562 cells were pulsed with YeiH.232-240 as described above and co-cultured with isolated CD8+ T cells from two participants with HLA-B*27-associated disease: one participant with both axSpA and AAU and one participant with only AAU.

After initial antigen stimulation, CD8+ T cells were cultured and expanded for 11 days. On day 11 of the culture, new YeiH-pulsed K562 cells were added to re-stimulate the expanded, antigen-specific CD8+ T cells. A custom Luminex multiplex assay was performed on the media supernatant to quantify the cytokine and chemokine output.

Flow cytometry was performed to determine the frequency of  YeiH+ CD8+T cells.

*Luminex Multiplex Cytokine Panel:*

A Luminex Multiplex Assay was run on media collected from the cultured CD8+ T cells before and after re-stimulation. The assay tested for the following 27 molecules in media: MIP-1 alpha, IL-27, IL-1 beta, IL-2, IL-4, IL-5, IP-10, IL-6, CD137, IL-8, IL-10, IL-13, IL017A, RANTES, IFN gamma, GM-CSF, TNF alpha, MIP-1 beta, IFN alpha, MCP-1, IL-9, perforin, IL-12/IL-23 p40, IL-18. Granzyme B, Granzyme A, and IL-22. Data was analyzed with Belysa 1.2.1 and plotted using GraphPad PRISM (v11.0.2) and R (4.4.0).
